# Role of Long-range Allosteric Communication in Determining the Stability and Disassembly of SARS-COV-2 in Complex with ACE2

**DOI:** 10.1101/2020.11.30.405340

**Authors:** Mauro L. Mugnai, Clark Templeton, Ron Elber, D. Thirumalai

## Abstract

Severe acute respiratory syndrome (SARS) and novel coronavirus disease (COVID-19) are caused by two closely related beta-coronaviruses, SARS-CoV and SARS-CoV-2, respectively. The envelopes surrounding these viruses are decorated with spike proteins, whose receptor binding domains (RBDs) initiate invasion by binding to the human angiotensin-converting enzyme 2 (ACE2). Subtle changes at the interface with ACE2 seem to be responsible for the enhanced affinity for the receptor of the SARS-CoV-2 RBD compared to SARS-CoV RBD. Here, we use Elastic Network Models (ENMs) to study the response of the viral RBDs and ACE2 upon dissassembly of the complexes. We identify a dominant detachment mode, in which the RBD rotates away from the surface of ACE2, while the receptor undergoes a conformational transition which stretches the active-site cleft. Using the Structural Perturbation Method, we determine the network of residues, referred to as the Allostery Wiring Diagram (AWD), which drives the large-scale motion activated by the detachment of the complex. The AWD for SARS-CoV and SARS-CoV-2 are remarkably similar, showing a network that spans the interface of the complex and reaches the active site of ACE2, thus establishing an allosteric connection between RBD binding and receptor catalytic function. Informed in part by the AWD, we used Molecular Dynamics simulations to probe the effect of interfacial mutations in which SARS-CoV-2 residues are replaced by their SARS-CoV counterparts. We focused on a conserved glycine (G502 in SARS-CoV-2, G488 in SARS-CoV) because it belongs to a region that initiates the dissociation of the complex along the dominant detachment mode, and is prominent in the AWD. Molecular Dynamics simulations of SARS-CoV-2 wild-type and G502P mutant show that the affinity for the human receptor of the mutant is drastically diminished. Our results suggest that in addition to residues that are in direct contact with the interface those involved in long range allosteric communication are also a determinant of the stability of the RBD-ACE2 complex.

## Introduction

Due to the unprecedented toll that COVID-19 has taken throughout the world, the extraordinary interest elicited by this pandemic from the scientific, medical, and pharmaceutical communities as well as the general public cannot be overstated. The causative virus in the COVID-19 disease is a novel beta-coronavirus. The membrane of this virus is decorated with homotrimeric spike proteins, which are directly responsible for initiating host invasion. The tip of the spike protein fluctuates between “down” and “up” conformations.^1^ In the “up” state, a receptor binding domain (RBD) is exposed to solution, and is poised to attach to the host, the first step of viral invasion. Experiments^2,3^ have established that the presence of the angiotensin converting enzyme 2 (ACE2) receptor is necessary for infecting the human cell. In addition, structural studies revealed the conformation of the full spike,^1,4^ of the RBD-ACE2 complex,^5-7^ and of the receptor-bound spike,^8^ thereby setting the stage for getting near quantitative molecular insights using computational methods.^9,10^

Nearly 20 years ago, the severe acute respiratory syndrome (SARS) raised worldwide health concerns. This disease was caused by another beta-coronavirus, SARS-CoV. Similar to the novel coronavirus (which from now on will be termed SARS-CoV-2), SARS-CoV invades the human cell by binding to the ACE2 receptor.^11,12^ Structure^4^ and sequence^2^ of SARS-CoV-2 and SARS-CoV spike proteins are similar (see Fig. 1 for a comparison of the RBDs), although the RBD of SARS-CoV-2 has higher affinity for ACE2.^4,7,13-15^ Binding assays and careful inspection of the RBD-ACE2 interface inform us of the relevant differences between the two viruses.^16,17^ Deep Mutagenesis Scanning provides the methodology for more extensive analysis. This technique enables a rapid evaluation of the effect on expression and binding of replacing selected ACE2^18^ and SARS-CoV-2 RBD^15^ residues with any other amino acid. This wealth of information is crucial in order to understand the mechanism by which RBD recognizes the ACE2. However, it needs to be complemented with detailed computational models capable of providing a structural rationale to the experimental data.

**Figure 1:**
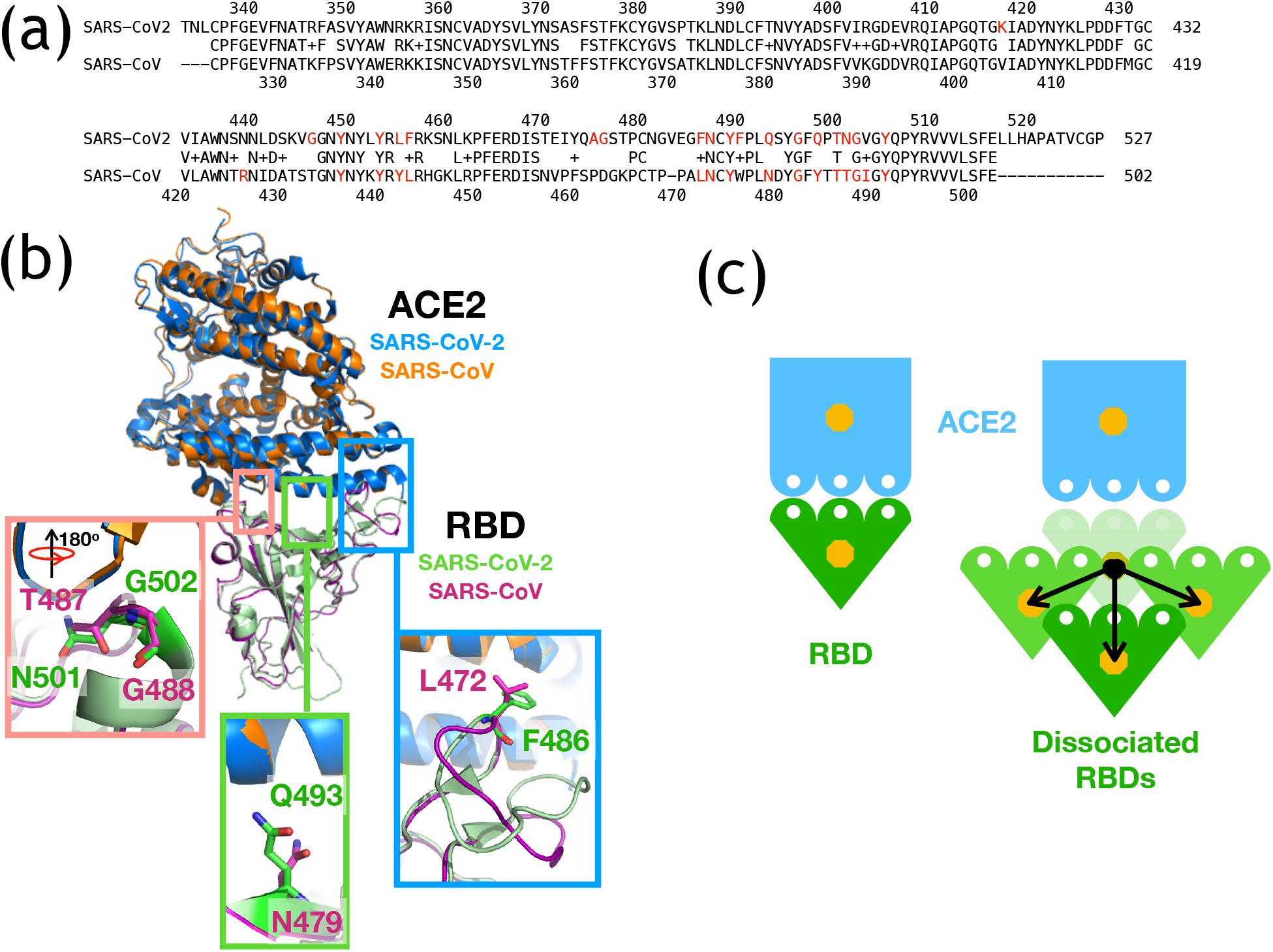
RBD-ACE2 complex. (a) Sequence alignment of SARS-CoV and SARS-CoV-2 RBDs. The sequences were aligned as described in the Methods section. Red residues engage in direct interactions with the interface of ACE2, that is any of their heavy atoms within 4 Å from a heavy atom of ACE2. (b) Comparison of the structures of the RBD-ACE2 complex for SARS-CoV (PDBID: 2AJF^25^) and SARS-CoV-2 (PDBID: 6LZG^6^). The SARS-CoV-2 complex is shown in blue (ACE2) and green (RBD), whereas the ACE2 and RBD of SARS-CoV are in orange and magenta, respectively. Only the ACE2 proteins were aligned using PyMol. ^26^ The boxes highlight three regions of interaction at the interface between the RBD and ACE2. The highlight in the red box is rotated by about 180° compared to the main figure. (c) Pictorial representation of the associated (left) and dissociated (right) complex used in the SPM analysis (see Supporting Information for details outlining the pathways of disassembly).

Binding affinity changes upon mutation could result from perturbation of local interactions (e.g. complementarity of charged, and hydrophobic patches), and long-range effects, which are often associated with large structural transitions.^19^ Such an “action at a distance” is encoded in the sequence, conformation, and dynamics of the protein or the associated complex. In the present context, the allosteric “down” ⇌“up” transition of the spike protein, as well as the binding/dissociation of the RBD and ACE2, likely involve a network of residues throughout the ensemble, as in other large information-transmitting complexes. Computational methods have been developed to predict long-range effects associated with such structural transitions, and provide complementary information to the analysis based solely on structure and sequence. The Structural Perturbation Method (SPM)^20-22^ enables the determination of the Allostery Wiring Diagram (AWD), which represents a network of residues that drives large-scale mechanical movement. SPM analysis begins with an Elastic Network Model (ENM)^23,24^ representation of the complex. The ENM is a coarse-grained description of the system in which all of the components interact via harmonic potentials. This simplified energy function underlies the fluctuations of the system, which are described using a combination of normal modes. The AWD determination using the SPM is based on the premise that key residues associated with conformational transitions between distinct allosteric states have the largest effect on the few low-frequency, soft modes that describe the large-scale movement. This can be probed by perturbing the interactions of each residue and monitoring the response along a mode. The residues that have the largest response to the perturbation are considered “hot-spots,” and the network of such residues is the AWD.

Here, we used the SPM analysis based on the structures of the complexes between the RBDs of SARS-CoV^25^ and SARS-CoV-2^5,6^ with the soluble, metallopeptidase domain (PD) of the human ACE2 receptor (Fig. 1b). ENM analysis revealed that a single dominant mode, which is similar in SARS-CoV and SARS-CoV-2 (see Fig. 2 and Fig. S1), describes the disruption of the complex. Along this mode, a subset of interfacial interactions focused around RBD residue G502 (G488 in SARS-CoV) appear to rupture first as the RBD undergoes a counter-clockwise rotation to move away from ACE2 (Fig. 2 and Fig. S1). The SPM analysis suggests an allosteric connection between the active site of ACE2 and the RBD-binding surface of the receptor (Fig. 3 and Fig. S2). At the interface between ACE2 and RBD, SPM identifies three distinct regions of interaction, which span the whole interface (Fig. 4 and Fig. S4). We identified a few residues that could contribute to the change in the affinity for the receptor of SARS-CoV and SARS-CoV-2 RBDs, which prompted us to perform Molecular Dynamics (MD) simulations to assess their effect on the stability of the complex and provide atomically-detailed explanation of our findings. SPM also predicts that the strongest contribution to the AWD comes from the residues around SARS-CoV-2 G502 and SARS-CoV G488. We surmise that G502 might play a critical role in the dissociation of the complex, and we tested this hypothesis by performing MD simulations of the G502P mutant. As predicted by the SPM, the mutation strongly destabilizes the complex. Taken together, our results show that disassembly of the RBD-ACE2 complex, and hence its stability, is determined by allosteric communication between residues that are spatially distant from the interface. A clear implication is that long range allosteric transitions must play a crucial role in driving the “down” ⇌“up” transition in the spike protein, which is the first step in viral infection.

**Figure 2:**
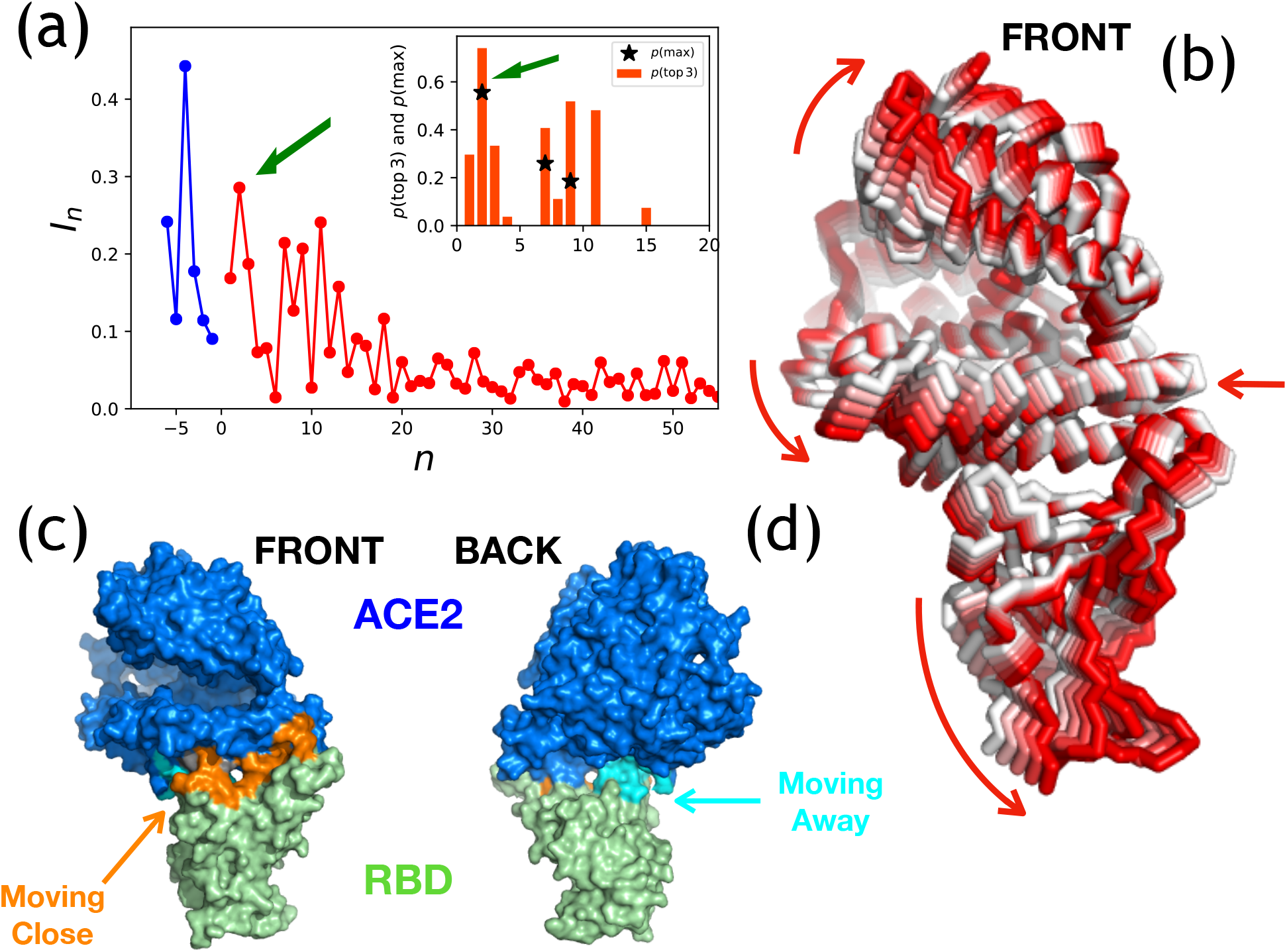
ENM analysis for ACE2-SARS-CoV-2 (PDBID: 6LZG^6^). (a) Overlap [Eq. (S18) in the Supporting Information] between the displacement vector associated with the transition from associated to dissociated complex (Fig. 1d). The overlap between the ensemble of displacement vectors and the first ≈ 50 eigenvectors, sorted in ascending order of the eigenvalues, as a function of the index of the eigenvectors. The first 6 eigenvectors, corresponding to the rigid body movement of the complex, are labeled with negative index and shown in blue. They are excluded from the rest of the analysis. The inset shows the probability within the set of 27 displacements considered that each mode is the top mode (black star) or is in the top 3 (red bar). The dominant mode, mode 2, is highlighted by a green arrow. (b) Movement associated with mode 2. The color scale (white-to-red) shows the direction of movement. The arrows highlight the displacement. (c-d) Highlight of mode 2 at the interface between ACE2 (blue) and RBD (green), with a view of the complex from the “front” (c) and “back” (d, rotated by ≈ 180°). Residues colored in cyan (orange) identify an area of the complex in which ACE2 and RBD are separating (getting closer) along mode 2. We computed the relative displacement between two beads at the complex interface (one ACE2, one RBD, within 8 Å from each other), that is 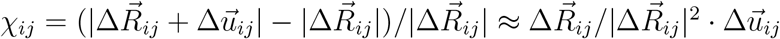. The distribution of *X_ij_* has nearly zero average and standard deviation *σ_x_*. If a residue *i* is such that *X_ij_* < —*σ_x_* for at least one *j*, and *X_ij_* < *σ_x_* for all *j*-s, we color that residue in orange. In contrast, we color a residue in cyan if *χ_ij_* > *σ_x_* for at least one *j*, and *χ_ij_* > —*σ_x_* for all *j*-s.

**Figure 3:**
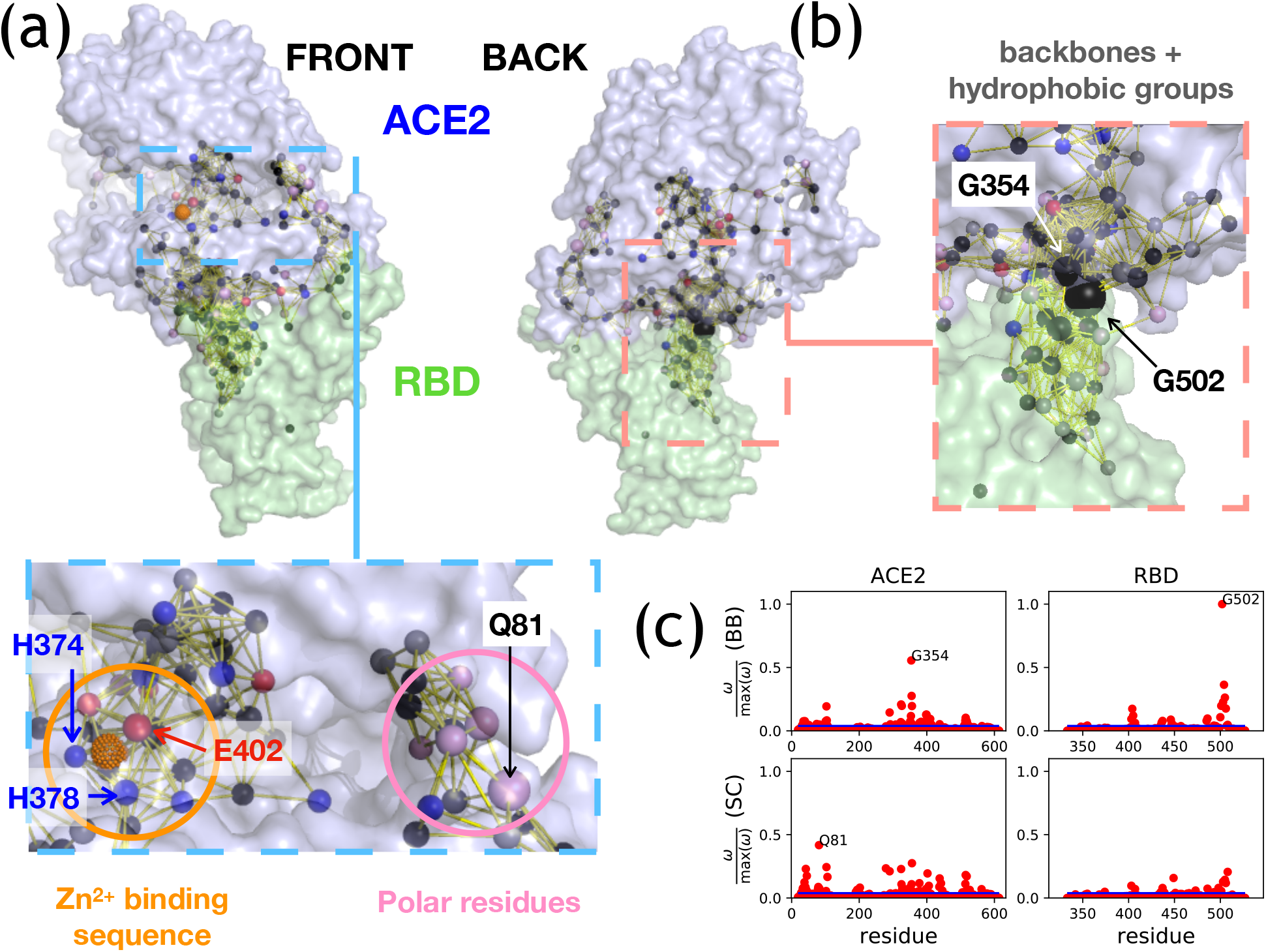
AWD for SARS-CoV-2. (a-b) The semi-transparent surface shows the 2.50 Å X-ray structure of SARS-CoV-2 complex (PDBID: 6LZG^6^). The human ACE2 is in blue and the viral RBD in green. The same complex is shown from the “front” (a) and the “back” (b) rotated by ≈ 180°. The spheres represent the beads selected by SPM as “hot spots”. The color code distinguishes the alpha-carbons (black), and side-chains which are positively charged (blue, including histidine), negatively charged (red), polar (pink), and hydrophobic (grey). The size of the sphere is α 0.5 + *ω_i_*/max *ω*, where *ω_i_* is the strength of the SPM signal for bead *i*, and max *ω* is the largest SPM signal. Beads selected by SPM and within 0.8 nm from each other are connected by a yellow line. (a) In the blue, dashed box, we highlight two sections of the AWD, which connect the bulk of the ACE2 with the interface. On one side (pink circle), communication is carried out through a cluster of predominantly polar side-chains, including Q81. On the other (orange circle), it reaches the area surrounding the zinc-binding consensus sequence, ^374^HExxH^378^+E^402^. (b) The red, dashed box from the back view shows a large cluster of predominantly alpha-carbons and hydrophobic side-chains that connect the interface of the complex with the rest of the ACE2 and RBD. The zoom-in box (red, dashed) indicates the presence of two glycine carrying a strong SPM signal: G354 on ACE2, and G502 on the RBD. (c) Ratio between the SPM signal of all the beads and the maximum signal. The top panels refer to backbone beads, the bottom two indicate sidechains. The left panels show the results for ACE2, the RBD is reported in the right panels. The blue line indicates the SPM cutoff (see Eq. S25).

**Figure 4:**
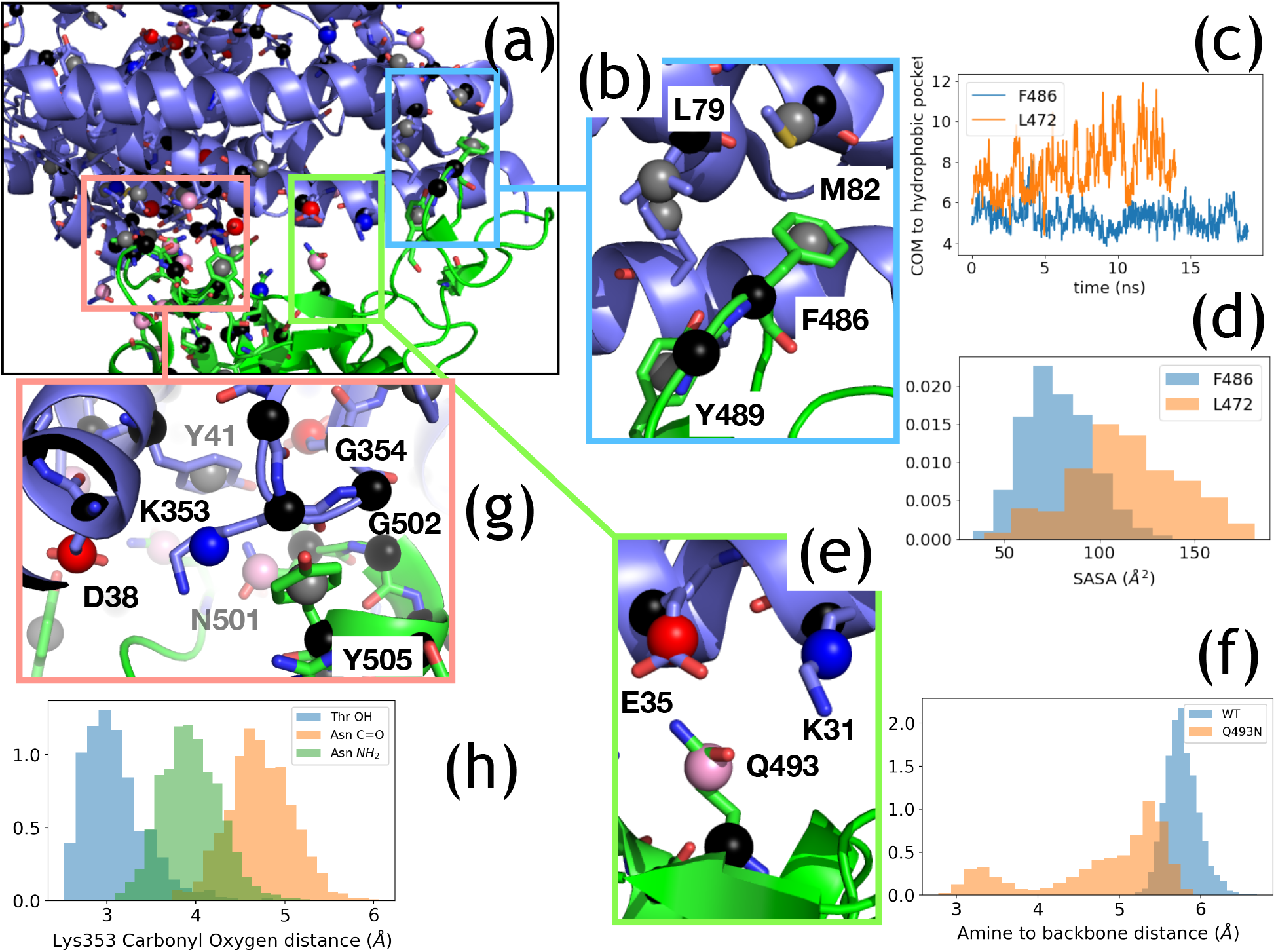
AWD at the interface for the SARS-CoV-2 complex (PDBID: 6LZG^6^). (a) Interface of the complex showing ACE2 (blue) and the RBD (green) in cartoon with side chains of “hot-spot” residues in stick-mode. The spheres indicate the beads determined by the SPM; the color of the bead is black for alpha-carbons, and for side-chains blue, red, pink, and grey indicate bases, acids, polar, and hydrophobic groups, respectively. (b) Hydrophobic cluster at the interface, showing contacts between F486 of RBD and M82 and L79 of ACE2. (c-d) MD simulations showing that F486 fits better than a leucine (the corresponding residue in SARS-CoV) in this hydrophobic pocket. (c) Distance between the RBD residue (F in blue, and L in orange) and M82 on ACE2. (d) Solvent-accessible Surface Area (SASA) of F (blue) and L (orange). (e) Residues from the previously identified “hot spot” 31: K31 and E35 form a salt bridge and engage in interaction with Q493 from the RBD. (f) Distribution of the distance between the end of Q493 (blue, WT SARS-CoV-2) and N493 (orange, mutant SARS-CoV-2) on RBD from their backbone: Q493 remains more extended, whereas N493 recoils on itself. (g) The residues of the well-known “hot spot 353” are predicted by the SPM: the ACE2 salt bridge K353-D38 is selected, together with the hydrophobic groups (Y41 on ACE2, and Y505 on the RBD), the polar N501 (RBD), and two glycine residues, G502 on the RBD, and G354 on ACE2. (h) The probability distribution of the distance between the carbonyl oxygen on the backbone of K353 and the polar atoms on residue 501 for both the WT SARS-CoV-2 (N, green for the amine, orange for the carbonyl) and the SARS-CoV analog (T, blue).

In the main text, we present results for only SARS-CoV-2 (Figs. 2-4). The Supporting Information contains the same analysis conducted on SARS-CoV (Figs. S1-S4). In the Discussion section we examine our findings in the context of few other published studies.

## Results

### Sequence Comparison

We compared the RBD sequences (see Methods) corresponding to the experimentally solved structures of SARS-CoV-2 and SARS-CoV (Fig. 1a) used in this study (for example see also^7,14,27^). About 81% of the residues are identical or similar (with a positive score in the replacement matrix, see Methods) between SARS-CoV and SARS-CoV-2. Focusing on residues involved in interfacial contacts (residues in red in Fig. 1a) only about 58% are identical or similar. We highlight a few sites which we will describe in detail in the following. (i) F486 in SARS-CoV-2 replaces L472 in SARS-CoV (red box in Fig. 1b). These residues are at the interface with ACE2 in a region that is completely reshaped in SARS-CoV-2 compared to SARS-CoV. This is due to significant differences in sequence – the sequence segment from 470 to 485 has only 3 conserved residues with 457-471 of SARS-CoV. (ii) Q493 of SARS-CoV-2 substitutes N479 in SARS-CoV in the middle section of the ACE2-RBD interface (green box in Fig. 1b). (iii) On the backside of the ACE2-RBD interface, N501 of SARS-CoV-2 has replaced SARS-CoV T487 (blue box in Fig. 1b). (iv) G502 of SARS-CoV-2 is conserved (G488 in SARS-CoV, see blue box in Fig. 1b).

### Dominant Mode

The dynamics of the ACE2-RBD complex can be described by performing a normal mode analysis of the ENM for the complex. Often, long-range, collective movements are encoded within a small set of low-eigenvalue modes. Therefore, we sought the modes that provide the best description of the dissociation of the viral RBD and the receptor. Because the mechanism of complex disruption is unknown, we considered 27 dissociation pathways (see Fig. 1c for a pictorial representation, and for additional details see the Supporting Information (SI) and Fig. S6). We computed the overlap (the cosine of the angle between two vectors, see Eq. S18) between each normal mode and all of the displacement vectors representing the disassembly of the complex. We neglected rigid body movements (colored blue, with negative indices in Fig. 2a and Fig. S1a), which are not associated with the disruption of interactions at the interface. For both SARS-CoV-2 (Fig. 2a) and SARS-CoV (Fig. S1a), mode 2 (with the second smallest, non-zero eigenvalue) has the largest overlap when averaged over the 27 displacement vectors (green arrow in Fig. 2a and Fig. S1a, see Eq. S19). Mode 2 is also the dominant mode for > 50% of the displacement vectors considered (insets in Fig. 2a and Fig. S1a, black stars), and it is in the top three for > 60% of the dissociation pathways (insets of Fig. 2a and Fig. S1a, red bars). Although other modes could be relevant, we focused only on the dominant one.

Figure 2b shows the movement associated with mode 2: the “jaws” (subdomain I and subdomain II, see Towler *et al.*^28^) enclosing ACE2 active site^28^ open slightly, and the N-terminal helix moves right to left (see horizontal red arrow). The RBD rotates away from ACE2 in a counter-clockwise manner. Figures 2c-d show that the interacting pairs at the interface in the “front” of the complex (Fig. 2c) get closer, whereas there is domain visible from the “back” of the ensemble (Fig. 2d) that separates along the dominant mode. Figures S1b-d show that mode 2 of SARS-CoV RBD-ACE2 complex shares analogous features.

### SPM and global AWD

Figures 3-4 show the result of the SPM analysis for SARS-CoV-2. The results for SARS-CoV are in Figs. S2-S4. We focus on the “hot spot” residues that show the largest responses to a local perturbation (Eq. S20). The network of “hot spot” residues builds the AWD.

The AWD in Figs. 3a,b shows how the response to detachment involves more than merely the residues at the ACE2 interface. Two pathways connect the interior of ACE2 with the interface (Fig. 3a): one dominated by polar residues, including Q81 (pink circle in Fig. 3a), which has amongst the strongest SPM signals (Fig. 3c). The other pathway reaches the active site of ACE2 PD, with the zinc-binding consensus sequence (^374^HExxH^378^ + E^402^, see Towler *et al.* ^28^) involved in the network (zinc is shown in orange in Fig. 3a). From the back (Fig 3b), it is clear that the region associated with the initiation of the detachment (cyan in Fig. 2d) is dominated by backbone beads and hydrophobic side-chains. In this region, residues G502 on the RBD and G354 of ACE2 show strong SPM signals (Fig. 3c).

The shape of the overall AWD is similar for SARS-CoV-2 and SARS-CoV (see Fig. 3a,b with Fig. S2a,b). A quantitative comparison between the two AWDs shows a robust correlation (Pearson coefficient ≥ 0.8, see Fig. S3), although there are a number of outliers reflecting the reorganization of the network due to mutations and subtle structural changes. Many of these outliers occur around the area at the “back” of the complex which initiates detachment (see Fig. S3). Here, both G502 and V503 in SARS-CoV-2 and the corresponding SARS-CoV residues (G488 and I489) have among the strongest SPM signals. However, the contribution of the glycine is larger in SARS-CoV-2 than in SARS-CoV, whereas SARS-CoV I489 has a much stronger signal than the corresponding SARS-CoV-2 valine.

### AWD at the Interface

We draw a number of inferences by examining the three “hot spot” areas at the interface (Fig. 4). Where possible, the conjectures are tested using MD simulations.

(A) SPM identified a group of predominantly hydrophobic residues concentrated around M82 of ACE2 (blue box in Fig. 4a and Fig 4b for SARS-CoV-2, see Fig. S4a,b for SARS-CoV). L79 is also part of this hydrophobic patch in the AWD on ACE2 for both SARS-CoV and SARS-CoV-2. In contrast, Y83 contributes to the AWD only for SARS-CoV. In both viruses, a hydrophobic group interacts with M82 and L79: F486 in SARS-CoV-2 or L472 in SARS-CoV. Because phenylalanine is larger and more hydrophobic than leucine, it fits better in the pocket around M82, allowing us to suggest that the former interacts more tightly with ACE2, despite the reduced SPM signal for F486 compared to L472.

To confirm this hypothesis, we performed MD simulations of each complex separately and show that L472 slips out of the pocket within a few nanoseconds, whereas F486 remains inside the pocket (see Fig. 4c). An analysis of the Solvent-accessible Surface Area (SASA) for F486 and L472 provides the same insight (see Fig. 4d). We also computed the relative binding strength (ΔΔ*G*) between the wild-type (WT) F486 SARS-CoV-2 and the mutant F486L. In order to do this, we used alchemical methods^29^ (see Methods and Supporting Information) to compute the change in free energy upon mutation of the RBD in isolation or in complex with ACE2. This can be related to the ΔΔ*G* by constructing the thermodynamic cycle (see Fig. S8). The resulting ΔΔ*G* (see Table 1) suggests that the mutant has reduced affinity for ACE2.

**Table 1:**
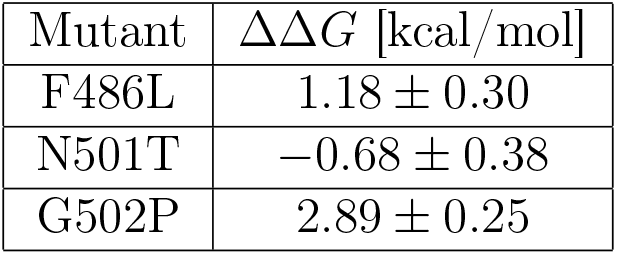
Results of alchemical substitutions. ΔΔ*G* are obtained averaging 3 independent runs, with error bars taken from the standard deviation. Positive ΔΔ*G* implies destabilization of the mutant.

(B) In the middle of the ACE2-RBD interface, SPM identified a second cluster around ACE2 residues K31 and E35 (green box in Fig. 4a and Fig 4e, see Fig. S4a,c for SARS-CoV), which form the previously identified “hot-spot 31” involving a salt bridge that could stabilize the complex.^16,17^ K31 and E35 interact with Q493 on the RBD of SARS-CoV-2, which is also a part of the AWD. In contrast, in SARS-CoV the Q493 is replaced by an asparagine (N479), and SPM displays fewer residues in this region of the AWD (see Fig. S4b).

We tested the difference between asparagine and glutamine by running MD simulations of the ACE2-RBD WT and Q493N mutant of SARS-CoV-2. Addition of another carbon in the side-chain allows Q493 to interact with K31 and E35 on the ACE2, whereas the shorter N493 does not appear to be able to engage in a stable interaction with the same residues on the ACE2 interface. As a consequence, Fig. 4f shows that N493 of the mutant, but not the WT Q493, spends a fraction of the time recoiled onto itself. Accordingly, in the ACE2-RBD complex of SARS-CoV, K31 and E35 move away from the RBD and interact with ACE2 Q76 (see PDBID: 2AJF^25^).

(C) On the left of the complex, there is a large cluster of residues focused around the salt-bridge between K353 and D38 on ACE2 (red box in Fig. 4a and Fig 4g, see Fig. S4a,d for SARS-CoV), which has been termed “hot spot 353”.^16,17^ Zooming in (Fig. 4g), we focus on a subset of residues identified by SPM: two hydrophobic groups (Y505 on RBD and Y41 on ACE2), G502 (RBD) and G354 (ACE2), which carry the largest SPM signals (see Fig. 3c) and N501 on the RBD. Similar groups are identified in SARS-CoV (Fig. S4d), though the SPM signal for K353 and D38 is weaker. Interestingly, N501 of SARS-CoV-2 replaces T487 in SARS-CoV. We tested the impact of this replacement by alchemically substituting N501 with a threonine, and we computed the relative binding free energy of ACE2 and RBD between the WT SARS-CoV-2 and the N501T mutant following the same thermodynamic cycle as for the F486L mutation (see Fig. S8). We found a negative ΔΔ*G* (see Table 1), implying that the mutant has a larger affinity for the human receptor, and we compared unbiased MD simulations of the WT and N501T mutant to determine the reasons of this change. From Fig. 4h, it appears that the hydroxyl group on the threonine can form a favorable hydrogen bond with the carbonyl backbone of K353. Given that the asparagine is slightly longer and only has a hydrogen bond donor at the end of its side-chain, it is unable to form this connection and is not long enough to successfully interact at the end of the K353.

At the core of the previously termed “hot-spot 353” is G502, which carries the largest SPM signal in SARS-CoV-2 and is in the top 3 for SARS-CoV. G502 is the proximal residue of a short near-alpha-helical turn made by an almost perfectly conserved set of 4 residues: ^502^GVGY^505^ and ^488^GIGY^491^ for SARS-CoV-2 and SARS-CoV, respectively. This turn on its own is unlikely to be stable in solution, given the lack of propensity for glycine to be in a alpha helix.^30^ MD simulations (Fig. 5a) confirm this showing that the RBD in solution is characterized by a larger average distance and broader distribution of the bond between G502 and Y505.

**Figure 5:**
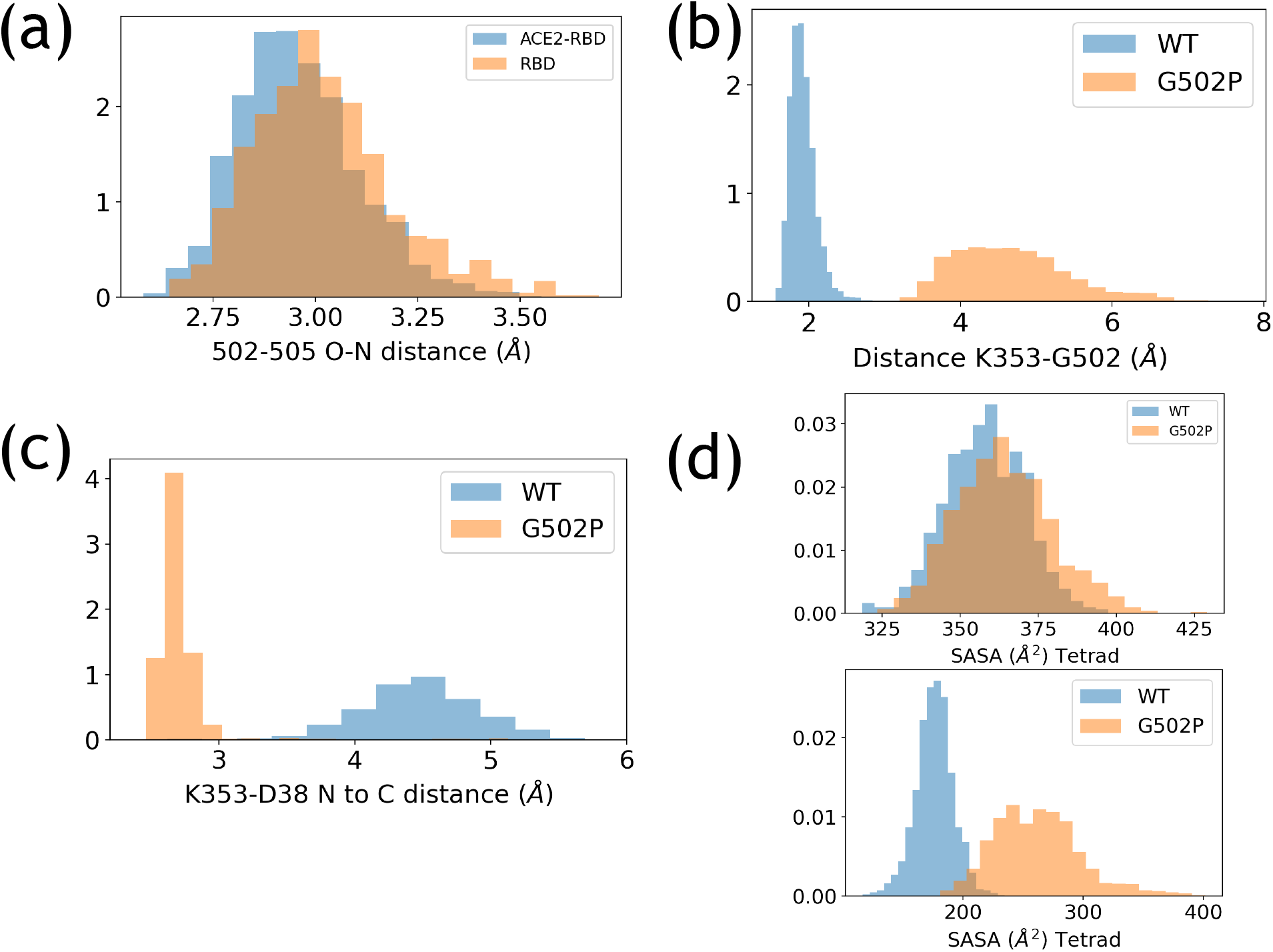
Structural impact of the G502 and the G502P mutation. (a) Distribution of the G502-Y505 backbone hydrogen bond for RBD in solution (orange) and in complex with ACE2 (blue). (b) Distribution of the distance between the backbone N of G502 (WT, blue) or P502 (mutant, orange) and the backbone carbonyl carbon of K353. (c) Distribution of the ACE2 D38-K353 distance for the WT (blue) and G502P (orange). (d) WT (blue) and mutant (orange) SASA distribution of the 502-505 tetrad for the solvated RBD (top) and the ACE2-RBD complex (bottom). The RBD shows little difference in solvent exposure upon mutation, while the complex indicates that the tetrad is more compact in the WT. Note that in the comparison between WT and G502P we account for the large size of proline by only considering the backbone of residue 502.

Importantly, G502 also forms a hydrogen bond with the backbone of K353, which likely stabilizes the complex. In addition, G502 tightly fits in a small recess on the surface of ACE2. ^15^ We mutated glycine 502 to proline, which has a dual effect: (i) it is incapable of interacting with K353, although it could in principle maintain the helical shape of the tetrad ^502^PVGY^505^, and (ii) it might introduce steric clashes with ACE2. The result indicates that proline strongly destabilizes the ACE2-RBD complex (see Table 1). Analysis of the MD simulations reveals that the distance between the RBD residue in position 502 and K353 increases after the mutation (Fig. 5b), an expected consequence of breaking the hydrogen bond. Interestingly, this is associated with the strengthening of the ACE2 K353-D38 saltbridge (Fig. 5c). The solvent exposure of the tetrad 502-505 (Fig. 5d) is unaffected by the mutation in the solvated RBD; on the other hand, residues 502-505 are more solvent exposed after the G502P mutation in the ACE2-RBD complex. This indicates that the increase in the volume from glycine to proline reduces the extent to which the tetrad 502-505 fits in the niche created by ACE2, which underscores the importance of volume-exclusion effects.

## Discussion

### Overall Movement of the Complex

Along the dominant mode, activated by the detachment, the interfacial contacts break in the vicinity of an area previously denoted as “hot spot 353.” An oscillating movement along the dissociation mode would predict the alternative weakening of the interaction at the two ends of the ACE2-RBD interface (the hydrophobic patch and “hot spot 353”), while the central area of the complex maintains its native distance from ACE2. Such oscillations along mode 2 would be in agreement with MD simulations of the complex, which show that the two ends of the interface break native contacts more frequently than the central section. ^31^

### The Importance of G502 (G488 in SARS-CoV)

The AWD probes the contribution of individual residues to the collective response associated with the dissassembly of the RBD-ACE2. However, the contribution of a residue to the AWD may not be directly correlated with its effect on the stability of the complex. This is clear from two considerations: first, a pair of interacting residues that do not move during the disassembly process would not contribute to the AWD, and yet might have a significant role in regulating the affinity of the complex – the interaction would remain intact, though it might be strengthened or weakened by changes in the environment. Second, the AWD only reveals information about the detachment, and not about the binding of RBD to ACE2. Nevertheless, the information from the AWD could be combined with other observations to predict the contribution of a specific group (backbone or side chain) to the stability of the complex. For example, a combination of factors suggest that modifying the interactions around the RBD G502 (G488 in SARS-CoV) could affect the stability of the complex. (1) SPM indicates that this glycine belongs to the part of the complex that appears to initiate detachment along the dominant mode (see Fig. 2d). (2) SPM also suggests that G502 is part of a branch of the AWD that reaches the bulk of ACE2 (Figs. 3a,b), highlighting that it contributes to a collective response, which is shown by its strong SPM signal (Fig. 3c). (3) Previous studies argued the importance of the K353-G502 hydrogen bond. ^32,33^ (4) Analysis of the structure suggests that the interaction with G502 might contribute to the stability of the ACE2 salt bridge between K353 and D38 on the receptor surface (Fig. 4g). This salt bridge could contribute to the stability of the complex because binding ACE2 to the RBD makes the environment more hydrophobic, which effectively reduces the dielectric constant making the K353-D38 salt bridge more stable than it would be in solution. ^17^ Indeed, structures of ACE2 without RBD show that the salt bridge could be either intact (PDBID: 6M18^5^) or broken (PDBID: 1R42^28^). (5) Computational studies indicated that residue G502 is replaced by proline in a number of coronaviruses which are predicted to interact poorly with the human ACE2 receptor.^34^ (6) Deep Mutagenesis Scanning indicated that mutation of G502 in any other residue hampers the ability of the RBD to bind the human receptor, and this is attributed to volume exclusion effects. ^15^

We provided structural characterization to explain these observations by studying the effect of the substitution of glycine 502 with proline using MD simulations. Free energy calculations showed that the mutant (G502P) complex is weakened by almost 3 kcal/mol (Table 1). We could ascribe this effect, at least in part, to the loss of the backbone hydrogen bond between G502 and K353. However, we also observed that the K353-D38 salt bridge is stabilized by this substitution. We surmise that the hydrogen bond with the RBD pulls K353 away from D38; in the absence of the interaction with the RBD, the lysine can move closer to the aspartic acid. While these opposing effects might balance each other, our calculations of the SASA for the the 502-505 tetrad show that the mutant does not fit well on the ACE2 surface, indicating the important role of steric clashes and shape complementarity. The importance of volume exclusion is highlighted in Fig. S9, where we plot the data of Starr *et al.*^15^ for the change in avidity of the mutant G502X as a function of the volume of the residue X.^35^

### Mutations at the Interface

We focused our analysis on a few amino acids at the interface which were replaced in SARS-CoV-2 compared to SARS-CoV. Our prediction of the change in binding affinity when F486 is mutated to leucine is in agreement with experiments measuring the change in binding avidity for the same mutation,^15^ and with experiments^16^ and simulations^33^ showing that the L472F mutation in SARS-CoV resulted in a more stable complex. Interestingly, the values for the ΔΔ*G* of F486L in SARS-CoV-2 and L472F in SARS-CoV^33^ are in excellent agreement. In addition, computational studies report a ΔΔ*G* for the F486A mutation that is somewhat larger than our result for the F486L substitution. ^36^ This is expected: alanine is smaller and less hydrophobic than leucine, resulting in a further binding penalty when it faces the ACE2 hydrophobic pocket around M82.

We found that the N501T mutation in SARS-CoV-2, which restores the SARS-CoV residue, increases the binding affinity of the RBD for ACE2. This is in agreement with expectations^17^ but contradicts other computational studies.^34^ Experiments have shown that the N501T reduces the binding strength of the full spike protein,^7^ although measurements probing the result for the RBD alone showed a small increase in avidity upon mutation.^15^ Interestingly, the inverse mutation, T487N, in SARS-CoV was also predicted via MD simulations to be favorable, ^33^ which might highlight that changes to nearby or far-away groups contribute to selecting which amino acid increases stability.

Our analysis of the difference between glutamine and asparagine in position 493 of SARS-CoV-2 indicates that there is a specific recognition of the RBD interface only with glutamine, which would indicate a small increase in binding affinity. We attempted to compute the ΔΔ*G* associated with this mutation, but preliminary results were deemed to be too small to be trustworthy. Deep Scanning Mutagenesis indicates that the Q493N mutation reduces the avidity of the ACE2-RBD complex by a small amount, though it also indicates that the specific recognition of K31 and E35 might not be crucial, as mutations to hydrophobic residues (M, A, or Y) further increase the avidity.^15^ Also in this case, the Q493N mutation reduces the binding affinity of the full spike for ACE2. ^7^ The inverse mutation on SARS-CoV, N479Q, was predicted to be favorable. ^33^

### Connection with the Active Site

A striking prediction of the SPM is the allosteric connection between the detachment of the RBD and the active site of ACE2 PD. The opening movement of the “jaws” of ACE2 (subdomain I and subdomain II) is opposite to the movement observed in the presence of an inhibitor of the catalytic function of the enzyme, ^28^ which instead triggers the closing of the two subdomains. In addition, the AWD triggered by the release of the RBD reaches the zinc-binding active site of ACE2. This evidence suggests a possible connection between enzymatic activity and the formation of the RBD-ACE2 complex. However, the experimental evidence obtained so far yielded mixed results. (i) Mutating the histidines of the zinc-binding consensus sequence to asparagines impedes catalysis, but did not prevent ACE2-RBD binding in SARS-CoV. ^11^ Similarly, (ii) mutations of H374 and H378 of ACE2 do not significantly enhance or deplete the frequency with which the ACE2 is selected by a cell expressing the SARS-CoV-2 spike compared to the wild-type. ^18^ On the other hand, (iii) an inhibitor of ACE2 was shown to reduce cell-fusion mediated by SARS-CoV. ^37^ A more direct test of the relationship between RBD binding and ACE2 inhibition is lacking, to our knowledge. The AWD would predict that a long-range connection between enzymatic active site and the RBD binding site exists, and although we cannot predict what the consequences would be, it indicates that further experiments are necessary to probe this allosteric effect.

### Importance of AWD

The presence of the AWD dispersed throughout the complex suggests that, besides the impact of electrostatic forces, ^38^ long-range effects caused by the inherent stiffness of the folded structure^19^ may be responsible in not only in the needed conformational transition that populates the infection competent “up” state, but also affect the responses at the interface. Although it may be tempting to focus solely on the interactions at the interface to infer stability of the complexes, residues that are spatially distant also affect binding interactions, and thus the efficacy of fusion. Thus, both for global mechanical movements through allosteric communication and for stability, it is critical to determine the AWD. ^21^

We focused our analysis on the RBD and the PD of ACE2, but excluded the role of sugars, which are known to be covalently linked to both ACE2 and RBD. The importance of glycosylation of the receptor in complex stability has already been investigated experimentally, and using MD simulations. ^7,18,31,39^ Future developments will expand the model of the viral protein including the rest of the spike, and of ACE2 (though the structure of the collectrin homology domain might not be stable in solution, as evidenced by the weak electron density map reported by Towler *et al.*,^28^ and therefore models that account for the cellular membrane might be required). The current study provides a sound computational framework to investigate the AWD activated transitions upon dissolution of the complex. Our study also suggests that allosteric communication across the entire spike may be involved in the initial stages of fusion.

## Materials and Methods

### Sequence Alignment

In order to compare the structures of the ACE2-RBD complexes (Fig. 1b), we aligned the sequences using dynamic programming.^40^ We adopted the BLO-SUM50 scoring matrix, with penalties −12 and —2 for opening and continuing gaps, respectively. ^40^ Aligned residues are considered similar if they have a positive BLOSUM50 score.

### ENM and SPM

The steps in the AWD determination are the following. First, we computed the modes and frequencies of the bound complexes (Fig. 1b), considering only protein components – RBD [333-527 for SARS-CoV-2 (PDBID: 6LZG), 323-502 for SARS-CoV (PDBID: 2AJF)] and ACE2 [19-614 for the complex with SARS-CoV-2 (PDBID: 6LZG), 19-615 for the complex with SARS-CoV) (PDBID: 2AJF)], ignoring ions (zinc and chloride, which is found in structure PDBID: 2AJF, but not in PDBID: 6LZG), water, and sugars. Second, we validated the model by comparing the scaled and normalized B-factors with experiments (see the Supporting Information and Fig. S5). The location and height of many peaks was reproduced correctly without adjusting any parameters. Third, we constructed an ensemble of dissociated structures (Fig. 1c, see details in the Supporting Information and Fig. S6) in which the ACE2 structure is intact, and the RBD is rigidly displaced away from the ACE2. Fourth, we identified the mode that provides the most accurate description of the rigid-body displacement of the RBD (one with the largest “overlap” defined in the Supporting Information, Eqs. S18-S19), and analyzed the mode-induced conformational changes in the ACE2 and RBD (Fig. 2 and Fig. S1). Finally, by performing the SPM analysis, we determine the critical residues throughout the complex (Fig. 3 and Fig. S2), and at the interface (Fig. 4 and Fig. S4). The SPM signal (*ω*, defined in Eq. S20) is a result of the response to a perturbation of the interactions of one group with the rest of the protein along a selected mode. The network of residues with the largest responses to a local perturbation (that is, 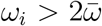, where 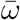 is the average SPM signal (see Eq. S25) constitute the AWD. Additional details of the ENM and SPM are in the Supporting Information.

### MD Simulations

We used alchemical substitutions^29^ in order to estimate the difference in binding free energy of the RBD-ACE2 complex upon mutation of a few RBD residues. The free energy change upon mutating a residue in the bound and dissociated complex can be computed numerically, and the scheme in Fig. S8 shows that it is related to the change in binding free energy (ΔΔ*G*) between the wild type and mutant. ΔΔ*G* > 0 refers to a mutation that decreases the stability of the complex, while ΔΔ*G* < 0 implies that the mutation stabilizes the ACE2-RBD complex.

The details of the MD simulations are discussed in the Supporting Information. Briefly, we performed the explicit-solvent simulations with NAMD^41^ with CHARMM36^42^ force field for proteins and TIP3P ^43^ for water. The VMD ^44^ package Mutator was used to setup the alchemical substitutions for computing the Δ*G* of mutation. The transformation of the WT protein into a mutant was carried out via a series of intermediate steps for both the RBD alone in solution, and the solvated RBD-ACE2 complex. The free energy difference was computed using Bennett Acceptance Ratio^45^ at each intermediate step. Results were repeated 3 times from different initial structures in order to estimate the error.

### Visualization

Molecular structures were rendered with PyMol^26^ or VMD.^44^ Matplotlib^46^ and Jupyter notebooks^47^ were also used to plot and analyze the data.

## Acknowledgement

We thank Jason McLellan for his interest, and Carlos Simmerling for useful comments. MLM thanks Atreya Dey for pointing out a typo in the Supporting Information. This work was supported by National Science Foundation (CHE 19-00093), National Institute of Health (NIH GM59796), the Welch Foundation F-1896, and F-0019 administered through the Collie-Welch chair.

## Notes

### Competing Interest Statement

The authors have declared no competing interest.

