## Supplementary Information for "Role of Long-range Allosteric Communication in Determining the Stability and Disassembly of SARS-COV-2 in Complex with ACE2"

#### Elastic Network Model

We construct Elastic Network Models (ENMs)[S1] from a second order Taylor series of the energy function based on the Self-organized Polymer model with Side-chains (SOP-SC).[S2] This formulation differs from the standard approach[S3] of constructing ENMs. Therefore, it is explained in detail here.

**SOP-SC model:** In the SOP-SC model, each residue is represented by two beads: a backbone bead, located on the alpha carbon, and a side-chain bead, at the geometric center of the resolved heavy atom of the side chain. We ignore the side-chain of glycine, which is the only residue represented by a single bead. The energy function is,

$$E_{\mathrm{SOP-SC}} = E_{\mathrm{bond}} + E_{\mathrm{native}} + E_{\mathrm{non-native}}. \tag{S1}$$

Bonded interactions are between backbone beads of residues that are consecutive in the chain, and between the backbone and side-chain bead of a specific amino acid. Let the position of the bead $i$ be $\vec{r}_i = \vec{R}_i + \vec{u}_i$, where $\vec{R}_i$ and $\vec{u}_i$ are the native-structure location and a (small) displacement from $\vec{R}_i$ of bead $i$, respectively. In the SOP-SC model, $E_{\mathrm{bond}}$ is given by a Finitely Extensible Non-linear Elastic (FENE) potential, which has three parameters: bond strenght ($k$), a maximum extension ($R_0$), and an equilibrium distance between the beads in the native conformation $|\Delta\vec{R}_{ij}| = |\vec{R}_i - \vec{R}_j|$. The FENE interaction is given by,

$$E_{\mathrm{bond},ij} = -\frac{1}{2}kR_0^2 \ln\Big[1 - \Big(\frac{|\Delta\vec{r}_{ij}| - |\Delta\vec{R}_{ij}|}{R_0}\Big)^2\Big], \tag{S2}$$

with $\Delta\vec{r}_{ij} = \vec{r}_i - \vec{r}_j$.

Native interactions are described by Lennard-Jones potentials and involve two residues that are at least three amino acids away along the sequence and are spatially within $R_{\mathrm{cut}} = 8$ Å from each other. Each interaction has two parameters: (i) the location of the minimum, which is given by the distance between the beads in the native structure, and (ii) the strength of interaction, which depends on the pair of beads considered,

$$E_{\mathrm{native},ij} = \epsilon_{ij}\Big[\Big(\frac{|\Delta\vec{R}_{ij}|}{|\Delta\vec{r}_{ij}|}\Big)^{12} - 2\Big(\frac{|\Delta\vec{R}_{ij}|}{|\Delta\vec{r}_{ij}|}\Big)^{6}\Big] \tag{S3}$$

When two backbone beads interact, the energy strength is given by $\epsilon_{BB}$; the parameter for interaction between a side-chain and a backbone bead is $\epsilon_{BS}$; if a contact occurs between two side-chain beads, the energy strength is given by $\epsilon_{SS}|\mathrm{BT}_{ij} - 0.7|$, where $\mathrm{BT}_{ij}$ is the parameter in the Betancourt-Thirumalai[S4] statistical potential associated with the residue pair $i$ and $j$. Non-native interactions occur between beads that are non-bonded and within

$\leq 2$ residues from each other, or at a distance larger than $R_{\text{cut}}$ in the native conformation. These are purely repulsive, and are given by,

$$E_{\text{non-native},ij} = \epsilon_I \left( \frac{\sigma_{ij}}{|\Delta \vec{r}_{ij}|} \right)^6, \tag{S4}$$

where $\epsilon_I$ is a new parameter, and $\sigma_{ij}$ is a measure of the size of the repulsive core for the interaction between beads $i$ and $j$. However, as explained in the next section, non-native interactions will be simplified to native-interactions in the context of the development of the Elastic Network Model.

**Elastic Network Model:** Elastic Network Models (ENMs) stem from the second order Taylor expansion of the energy function describing the interaction between the various components of a model. In our case it is based on the alpha-carbon and side-chain representation of the complex with the energy function given in Eq. S1. The energy associated with the displacement of the beads by $\vec{u}_i$ is:

$$E_{\text{SOP}} = \frac{1}{2} \sum_{i=1}^{N} \sum_{j=1}^{N} E_{ij}(|\Delta \vec{r}_{ij}|) = \frac{1}{2} \sum_{i=1}^{N} \sum_{j=1}^{N} E_{ij}(|\Delta \vec{R}_{ij} + \Delta \vec{u}_{ij}|), \tag{S5}$$

where $\Delta \vec{u}_{ij} = \vec{u}_i - \vec{u}_j$, and the energy function is assumed to be a sum of pairwise interactions depending only on the distance between two beads.

Under the assumption that $|\Delta \vec{R}_{ij} + \Delta \vec{u}_{ij}| \approx |\Delta \vec{R}_{ij}|$, we can expand each term of the energy function, $E_{ij}(|\Delta \vec{R}_{ij} + \Delta \vec{u}_{ij}|)$, around $\Delta \vec{R}_{ij}$; this leads us to,

$$\begin{aligned} E_{\text{SOP}} \approx E_{\text{SOP},0} &+ \frac{1}{2} \sum_{i=1}^{N} \sum_{j=1}^{N} \left( \frac{\partial E_{ij}}{\partial |\Delta \vec{r}_{i,j}|} \right)_{|\Delta \vec{u}_{ij}|=0} (|\Delta \vec{R}_{ij} + \Delta \vec{u}_{ij}| - |\Delta \vec{R}_{ij}|) + \\ &+ \frac{1}{4} \sum_{i=1}^{N} \sum_{j=1}^{N} \left( \frac{\partial^2 E_{ij}}{\partial |\Delta \vec{r}_{i,j}|^2} \right)_{|\Delta \vec{u}_{ij}|=0} (|\Delta \vec{R}_{ij} + \Delta \vec{u}_{ij}| - |\Delta \vec{R}_{ij}|)^2. \end{aligned} \tag{S6}$$

The first term in the expansion is the energy when $\Delta \vec{u}_{ij} = 0$, which is a constant, and hence does not affect our results. Therefore, we ignore it. We need to compute the first and the second derivative of all the $E_{ij}$-s, which come from the SOP-SC energy function. For the bonded and native interactions the first derivative is zero, and the second derivatives are,

$$\Big(\frac{\partial^2 E_{\text{bonded},ij}}{\partial|\Delta\vec{r}_{ij}|^2}\Big)_{|\Delta\vec{u}_{ij}|=0} = k, \tag{S7}$$

$$\Big(\frac{\partial^2 E_{\text{native},ij}}{\partial|\Delta\vec{r}_{ij}|^2}\Big)_{|\Delta\vec{u}_{ij}|=0} = 72\frac{\epsilon_{ij}}{|\Delta\vec{R}_{ij}|^2}. \tag{S8}$$

For non-native interactions, the first derivative would be non-zero as well. In order to simplify the model, we modify the SOP-SC energy function and assign all the non-bonded interactions between beads such that $|\Delta\vec{R}_{ij}| < R_{\text{cut}}$ (and regardless of the distance along the sequence between $i$ and $j$, as long as they are not bonded) a native functional form. In this way, we are left with the following:

$$E_{\text{ENM}} = \frac{1}{4}\sum_{i=1}^{N}\sum_{j=1}^{N} K_{ij}(|\Delta\vec{R}_{ij} + \Delta\vec{u}_{ij}| - |\Delta\vec{R}_{ij}|)^2, \tag{S9}$$

with $K_{ij}$ given by,

$$K_{ij} = k \cdot B_{ij} + 72\frac{\epsilon_{ij}}{|\Delta\vec{R}_{ij}|^2} \cdot \Theta(R_{\text{cut}} - |\Delta\vec{R}_{ij}|) \cdot (1 - B_{ij}), \tag{S10}$$

with $B_{ij} = 1$ if the beads are bonded and zero otherwise, and $\Theta(R_{\text{cut}} - |\Delta\vec{R}_{ij}|)$ is the step function [$\Theta(x) = 1$ if $x > 0$ and zero otherwise]. The parameters in the model are provided in Table S1.

Table S1: Parameters for ENM implementation of SOP-SC model.

| | | |
|---|---|---|
| $\epsilon_{\text{BB}}$ | 0.55 kcal/mol | backbone-backbone interaction |
| $\epsilon_{\text{BS}}$ | 0.4 kcal/mol | backbone-side chain interaction |
| $\epsilon_{\text{SS}}$ | 0.3 kcal/mol | side chain-side chain interaction |
| $k$ | 20 kcal/(mol Å$^2$) | covalent bond |
| $R_{\text{cut}}$ | 8 Å | cutoff of native interactions |

Next, we can expand $|\Delta\vec{R}_{ij} + \Delta\vec{u}_{ij}|$ to first order in $|\Delta\vec{u}_{ij}|$, and we obtain,

$$E_{\mathrm{ENM}} = \frac{1}{2}\mathbf{u} \cdot \hat{\mathrm{H}} \cdot \mathbf{u}, \tag{S11}$$

where $\mathbf{u}$ is a $3N$ vector given by,

$$\mathbf{u}^{\intercal} = [(\vec{u}_1)_x, (\vec{u}_1)_y, (\vec{u}_1)_z, (\vec{u}_2)_x, (\vec{u}_2)_y, (\vec{u}_2)_z, \cdots, (\vec{u}_N)_x, (\vec{u}_N)_y, (\vec{u}_N)_z], \tag{S12}$$

with $(\vec{u}_i)_\alpha$ corresponding to the $\alpha \in \{x, y, z\}$ component of the displacement vector for the $i$-th bead. Moreover, $\hat{\mathrm{H}}$ is the Hessian matrix, with the following block matrix form,

$$\hat{\mathrm{H}} = \begin{pmatrix} \hat{H}_{11} & \hat{H}_{12} & \cdots & \hat{H}_{NN} \\ \hat{H}_{21} & \hat{H}_{22} & \cdots & \hat{H}_{2N} \\ . & . & \cdots & . \\ \hat{H}_{N1} & \hat{H}_{N2} & \cdots & \hat{H}_{NN}. \end{pmatrix} \tag{S13}$$

The components of the 3x3 matrices, $\hat{H}_{ij}$, are,

$$(\hat{H}_{ij})_{\alpha\beta} = \begin{cases} \sum_{l=1}^{N} K_{il}(\Delta\hat{R}_{il})_\alpha(\Delta\hat{R}_{il})_\beta & \text{if } i = j \\ -K_{ij}(\Delta\hat{R}_{ij})_\alpha(\Delta\hat{R}_{ij})_\beta & \text{if } i \neq j \end{cases} \tag{S14}$$

with $\alpha \in \{x, y, z\}$ and $\beta \in \{x, y, z\}$.

**Diagonalization of the Hessian:** From the diagonalization of the Hessian, we obtained the list of eigenvalues $h_l$ and related eigenvectors $\vec{v}_l$ sorted in ascending order of $h_l$. As explained in the main text, the 6 smallest eigenvalues (index $l = 0, 1, \cdots, 5$) are practically 0 ($< 10^{-12}$), due to the invariance of the Hessian matrix upon rigid body translations and rotations. It is easy to show that the Hessian matrix respects these physical constraints (see, for instance, ref.[S5]).

### B-factors

**Calculation of B-factors:** The B-factor of residue $i$ is defined as,[S3]

$$B_i = \frac{8\pi^2}{3}\langle \vec{u}_i^2 \rangle, \tag{S15}$$

where the symbol $\langle \cdots \rangle$ refers to the thermal average of the displacement vector ($\vec{u}_i$) of the i-th bead. The expression above may be rewritten as follows,[S3]

$$B_i = \frac{8\pi^2 k_B T}{3} \sum_{l>5} \frac{\sum_{\alpha \in \{x,y,z\}} (\vec{v}_l)^2_{i,\alpha}}{h_l}, \tag{S16}$$

where $(\vec{v}_l)_{i,\alpha}$ is the component of the $l$-th eigenvector associated with the displacement of the $i$-th bead in direction $\alpha$, and $k_B T$ is the thermal energy.

**Comparison with Experiments:** We compared the B-factors from the model with the experimental B-factors. Instead of comparing the absolute values of the peaks, we focused on their location and relative heights. Therefore, we compared,

$$S_i = \frac{B_i - \overline{B}}{\sigma(B)}, \tag{S17}$$

where $B_i$ is the B-factor of bead $i$, $\overline{B}$ is the average over all beads, and $\sigma(B)$ is the standard deviation among all the beads. We note here that the average and standard deviations are computed for the whole complex. In Fig. S5 we compare the results for 3 different experimentally determined structures of SARS-CoV-2, and for a single structure of SARS-CoV ACE2-RBD.

Clearly, the model reproduces the location and relative amplitude of most peaks. It is important to note that the model performs better against structures solved at higher resolution (Fig. S5a,d). This suggests that larger deviations from experiments in Figs. S5b,c could be ascribed, at least in part, to the lower resolution of the experimental structure. Other factors include the presence of other proteins and ACE2 domains (collectrin and transmembrane helix) in the lower resolution structure, which are not included in the calculation.

**Numerical Implementation and Technical Details:** The ENM and SPM analysis were performed using a "in-house" code written in Python3, which makes use of NumPy[S6,S7] (v.1.14.0 and v.1.17.2) and SciPy[S8] (v.1.0.0 and v.1.3.1) libraries to carry out linear algebra operations. Eigenvalues obtained with different version of the libraries were identical up to $\approx 10^{-15}$, and the eigenvectors associated with non-degenerate eigenvalues (that is, not the 6 related with rigid body movement) were identical up to $\approx 10^{-15}$ and an overall (irrelevant) sign. SPM signals for mode 2 obtained with different libraries were identical up to $\approx 10^{-13}$ (data were saved to this precision).

ENMs were constructed starting from the experimentally solved crystal structure ignoring ligands (zinc ion, water) as well as non-protein components (sugars). Missing residues were not modeled. If multiple conformations of a side-chain were available, the one listed as "A" was selected.

### Ensemble of Detached States and Overlap

**Ensemble of Detached States:** The detached state of the complex is modeled as a conformation in which the structure of the ACE2 and RBD are not changed, but the RBD is rigidly displaced away from ACE2. The question is, what is the direction of this rigid displacement? This may depend on which side of the RDB detaches first from ACE2. Because the answer to this question is unknown we constructed an ensemble of possible geometries of the detached state using the algorithm illustrated in Fig. S6. Fig. S6a shows a pictorial representation of ACE2 (blue) and the RBD (green). The interaction sites at the interface are shown as white circles, and the geometric centers of the RBD and ACE2 are reported as orange stars. Each site of the RBD located at the interface interacts with interfacial sites of the ACE2 that are within $R_{\text{cut}} = 8$ Å. We created a list of vectors comprising all the interaction pairs involving a group of RBD and one of ACE2 (Fig. S6b, orange and yellow arrows). Next, we averaged these vectors and displaced the geometric center of the RBD along the average vector away from ACE2; the length of this displacement is set to the cutoff of the interactions, $R_{\text{cut}}$ (Fig. S6c). We refer to the resulting detached ACE2-RBD complex as the "central" one (Fig. S6d, the reason of the name will become clear soon). Starting from the "central" RBD, we displaced its geometric center by $\vec{\delta} = R_{\text{cut}}(i, j, k)$, where $i$, $j$, and $k$, are indexes that can assume value $-1$, 0, or 1, for a total of 27 $\vec{\delta}$ vectors – Fig. S6e shows only 2. The resulting ensemble of detached conformations for the RBD comprises RBD-s that have been displaced by a variable amount. This is unnecessary, and therefore we move all the non-"central" RBD-s closer to the bound conformation of the RBD, so that each one of them is obtained via a displacement of $R_{\text{cut}}$ in 27 different directions (see Fig. S6f).

What does this scheme probe? As is clear from the Hessian, the interactions in the ENM model are "directional", meaning that movements orthogonal to $\Delta\vec{R}_{ij}$ do not change the energy of the interaction between beads $i$ and $j$. Therefore, by considering multiple directions of the displacement vector, we are probing the conformations in which different sets of interactions are "stressed" in the detachment process. As alluded to before, we do not know which ones are broken initially during detachment, and by considering 27 different pathways we sample a variety of possibilities.

The displacement vectors (black lines in Fig. S6f) are referred to as $\mathbf{D}_i$ and have $3N$ dimensions, with $N$ being the total number of beads, that is $\mathbf{D}_i = [(\mathbf{D}_i)_{1,x}, \cdots, (\mathbf{D}_i)_{N,z}]$, where $(\mathbf{D}_i)_{j,\alpha}$ is the entry associated with bead $j$, direction $\alpha \in \{x, y, z\}$. Note that, by construction $(\mathbf{D}_i)_{j,\alpha} = 0$ if $j$ belongs to ACE2. Because the displacement is rigid, all the $(\mathbf{D}_i)_{j,\alpha}$ are the same for all the beads belonging to the RBD.

**Overlap Calculation:** The overlap between eigenvector $n$ and the displacement $\mathbf{D}_i$ is,

$$I_n(\mathbf{D}_i) = \frac{|\sum_{j=1}^{N}\sum_{\alpha}(\mathbf{D}_i)_{j,\alpha}(\vec{v}_n)_{j,\alpha}|}{\sqrt{\sum_{j=1}^{N}\sum_{\alpha}(\mathbf{D}_i)_{j,\alpha}^2}\sqrt{\sum_{j=1}^{N}\sum_{\alpha}(\vec{v}_n)_{j,\alpha}^2}}, \tag{S18}$$

where $\alpha$ refers to the three Cartesian coordinates, and $(\vec{v}_n)_{i,\alpha}$ is the element of the n-th eigenvector related to bead $i$, coordinate $\alpha$. Note that ultimately only the direction of the displacement matters, and not its modulus. The average overlap for eigenvector n is,

$$\overline{I}_n = \frac{\sum_{i=1}^{27} I_n(\mathbf{D}_i)}{27}, \tag{S19}$$

where the sum is over all the different displacement vectors.

#### Structural Perturbation Method (SPM)

Once a mode, $m$, is selected, our goal is to understand how different beads of the complex contribute to that mode. The idea of the Structural Perturbation Method (SPM) is to answer this question by locally perturbing the interactions involving a particular bead $i$, and monitor how this affects the selected mode, $\mathbf{v}_m$. In order to assess the response of the system, we computed,

$$\omega_n(\mathbf{v}_m) = \frac{1}{2}\mathbf{v}_m \cdot \delta_n \hat{\mathrm{H}} \cdot \mathbf{v}_m, \tag{S20}$$

where the matrix $\delta_n\hat{\mathrm{H}}$ is the change in the Hessian obtained by removing from the network the links connecting bead "n" with the rest of the system, that is

$$\delta_n\hat{\mathrm{H}} = \hat{\mathrm{H}} - \hat{\mathrm{H}}(K_{ni} = 0, \forall i \in [1, \cdots, N]). \tag{S21}$$

The idea is pictorially represented in Fig. S7. From Eqs. S13-S14, it can be shown that,

$$\delta_n \hat{\mathrm{H}} =$$

$$= \begin{pmatrix} \delta_n\hat{H}_{11} & 0 & \cdots & 0 & 0 & \delta_n\hat{H}_{1n} & 0 & 0 & \cdots & 0 \\ 0 & \delta_n\hat{H}_{22} & \cdots & 0 & 0 & \delta_n\hat{H}_{2n} & 0 & 0 & \cdots & 0 \\ . & . & \cdots & . & . & . & . & \cdots & . & . \\ 0 & 0 & \cdots & 0 & \delta_n\hat{H}_{n-1n-1} & \delta_n\hat{H}_{n-1n} & 0 & 0 & \cdots & 0 \\ \delta_n\hat{H}_{n1} & \delta_n\hat{H}_{n2} & \cdots & \delta_n\hat{H}_{nn-2} & \delta_n\hat{H}_{nn-1} & \delta_n\hat{H}_{nn} & \delta_n\hat{H}_{n+1n} & \hat{H}_{n+2n} & \cdots & \delta_n\hat{H}_{nN} \\ 0 & 0 & \cdots & 0 & 0 & \delta_n\hat{H}_{n+1n} & \delta_n\hat{H}_{n+1n+1} & 0 & \cdots & 0 \\ . & . & \cdots & . & . & . & . & & .\cdots & . \\ 0 & 0 & \cdots & 0 & 0 & \delta_n\hat{H}_{Nn} & 0 & 0 & \cdots & \delta_n\hat{H}_{NN} \end{pmatrix} \tag{S22}$$

where $\delta_n\hat{H}_{ij}$-s are 3x3 matrices given by,

$$(\delta_n\hat{H}_{ij})_{\alpha,\beta} = \begin{cases} \sum_{l=1}^{N} K_{nl}(\Delta\hat{R}_{nl})_\alpha(\Delta\hat{R}_{nl})_\beta & \text{if } i = n, j = n; \\ -K_{nj}(\Delta\hat{R}_{nj})_\alpha(\Delta\hat{R}_{nj})_\beta & \text{if } i = n, j \neq n; \\ K_{nj}(\Delta\hat{R}_{nj})_\alpha(\Delta\hat{R}_{nj})_\beta & \text{if } i = j, j \neq n; \end{cases} \tag{S23}$$

Clearly, $\delta_n\hat{\mathrm{H}}$ is a sparse matrix, in which the only non-zero elements are along the diagonal, and along row and column $n$. The matrix multiplications in Eq. S20 are performed using SciPy[S8] libraries for sparse matrices.

There is an alternative way to formulate the problem; $\omega_n(\mathbf{v}_m)$ can be written as,

$$\omega_n(\mathbf{v}_m) = \alpha^{-2}\sum_{j=1}^{N}\delta_n K_{nj}(|\Delta\vec{R}_{ij} + \alpha\Delta(\mathbf{v}_m)_{ij}| - |\Delta\vec{R}_{ij}|)^2. \tag{S24}$$

Here, $\Delta(\mathbf{v}_m)_{ij}$ is the difference between eigenvectors entries for beads $i$ and $j$, and $\alpha$ is a small parameter that ensures that this relative displacement away from the native distance is small. Because the eigenvectors are normalized to 1, the order of magnitude for each term in the eigenvector is $\sim 1/(3N)^{1/2}$, which is likely small enough for a large system; therefore $\alpha$ can be set to 1.[S9] Following the same expansion carried out to arrive to Eq. S11, we obtain Eq. S20.

Finally, a bead $i$ is deemed to be a "hot spot" (strong SPM signal) for the movement associated with the selected eigenvector if,[S9]

$$\omega_n(\mathbf{v}_m) > 2\frac{\sum_{i=1}^{N}\omega_i(\mathbf{v}_m)}{N}. \tag{S25}$$

### Analysis of SARS-CoV

We present results for ENM results for ACE2-SARS-CoV RBD in Figs S1-S4, mirroring those presented in Figs. 2-4. The PDBID:2AJF[S10] is used for the complex. Comparison of the results in Figs S1-S4 and Figs. 2-4 allows us to draw the following conclusions. (i) Mode 2 remains the dominant mode for SARS-CoV-2 (Fig. S1), which is not surprising given the high structural similarity. (ii) The AWDs of SARS-CoV and SARS-CoV-2 are superficially similar (Fig. S2) but with subtle differences that could contribute to affect the stability of the complex (Fig. S3). A detailed comparison between the strengths of the SPM signals (Fig. 3c in the main text and Fig.S2c) shows that there is a higher density of residues with near-maximal SPM signal, although the correlation of the $\omega_i$ values are high (Pearson correlation coefficient $\geq 0.8$, see Fig. S3). Strikingly, in SARS-CoV-2 G502 carries the strongest signal, with weaker contributions from other groups, including V503. The corresponding SARS-CoV residues, G488 and I489, are also important although I489 carries the largest signal (Fig. S3). Even though the signal for G488 is not maximal, it is near the top, and the pathways reaching the bulk of ACE2 have similar features in SARS-CoV and SARS-CoV-2. (iii) The interface of SARS-CoV has the same three domains of interaction (Fig. S4). There is an extra hydrophobic group (Y83) identified by SPM in the hydrophobic patch around M82, and "hot spot 31" is weaker, with no contacts found, and with neither bead on K31 identified by SPM. In “hot spot 353” the salt bridge K353-D38 is not fully identified by the SPM (the backbone of D38 and the side chain of K353 are missing), but a number of predominantly hydrophobic side chains and backbone beads contribute to the AWD.

#### MD Simulations

MD simulations were performed in order to compute the change in the binding free energy upon mutation. and to provide structural and dynamical insights into the stability of the complex. Free energy differences were computed using the thermodynamic cycle shown in Fig. S8. The vertical transitions refer to binding of the wild type (WT) and the mutant protein, and can be measured experimentally. The horizontal arrows refer to the free energy changes upon mutations within the bound complex or the RBD alone. The corresponding free energies may be computed using alchemical methods[S11] (see Methods section of the main text). Because free energy is state function, the sum of the free energies along a closed path in Fig. S8 is = 0. This allows us to compute the change in the binding free energy upon mutation using simulations by exploiting the following equality,

$$\Delta\Delta G = \Delta G^{\mathrm{Bind}}_{\mathrm{Mut}} - \Delta G^{\mathrm{Bind}}_{\mathrm{WT}} = \Delta G^{\mathrm{Complex}}_{\mathrm{WT}\to\mathrm{Mut}} - \Delta G^{\mathrm{RBD}}_{\mathrm{WT}\to\mathrm{Mut}}. \tag{S26}$$

As a result, a positive $\Delta\Delta G$ (that is $\Delta G^{\mathrm{Bind}}_{\mathrm{Mut}} > \Delta G^{\mathrm{Bind}}_{\mathrm{WT}}$) indicates that the WT bound complex is more stable. Conversely, the mutation increases the strength of the dimer if $\Delta\Delta G < 0$.

**Setup:** In order to setup the MD simulations, we used a different PDB structure from the ENM, that is PDBID: 6M17,[S12] using only residues up to 729 for ACE2 (chain B) and the full RBD (chain E), neglecting crystallographically solved atoms that did not belong to the protein – i.e., the zinc ion. From there a topology file was built using the CHARMM36[S13] force field. The system was solvated with TIP3P[S14] water such that there is 15 Å padding in all the directions, and 150 mM NaCl was added using VMD plugins “autosolvate” and "autoionize."

**Equilibration:** Conjugate gradient energy minimization was performed for 3,000 steps followed by 500 ps in NVT and 500 ps in NPT to equilibrate the pressure and volume. During equilibration time, the atoms of the protein complex were kept constrained. Temperature was regulated at 310K with a Langevin piston of 10 $ps^{-1}$; electrostatics and Van der Waals (VdW) interactions were evaluated with a 12 Å cutoff. Long range electrostatics were computed using the Particle Mesh Ewald (PME) with a grid spacing of 1 Å. From this a 30 ns trajectory was generated, releasing the complex, prior to any data sampling or alchemical analysis in order to allow for additional equilibration. The center of mass of ACE2 was restrained using a harmonic spring with a force constant 10 kcal/(mol $Å^2$) to prevent drifting. All the simulations were performed using the software package NAMD.[S15]

**Alchemical Mutagenesis:** To perform the alchemical mutagenesis study, the VMD[S16] package "Mutator" was used to create a dual topology file. The free energy change can be evaluated using free energy perturbation techniques whereby the amino acid residue is slowly mutated according to a Hamiltonian that is dependent on a switching parameter $\lambda_{\mathrm{VdW}} \in [0, 1]$: $\lambda = 0$ corresponds to the WT, and $\lambda = 1$ to the mutant. The switching parameter was changed from 0 to 1 by taking 20 equally sized steps. Each lambda value was equilibrated for 100 ps before a data collection period of 1.5 ns with a sampling rate of the energy difference every 2 ps. For the G502P mutation, we increased the data collection period to 3 ns as this mutation provides a stronger perturbation to the structure. In order to avoid the end-point catastrophe, the van der Waals (VdW) interactions were treated using a soft-core potential,

$$r^2 = r^2 + \begin{cases} 6.5\lambda & \text{for a WT atom} \\ 6.5(1-\lambda) & \text{for a mutant atom} \end{cases}$$

Mutant (WT) electrostatic interactions were switched on (off) for $\lambda \in [0.5, 1]$ ($\lambda \in [0, 0.5]$) in order to ensure that the repulsive core (VdW interactions) was strong while atoms were charged. This protocol was determined by setting the parameter “alchElecLambdaStart” to 0.5.

To estimate the free energy difference for a single lambda value, we performed simulations in both the forward and backward direction such that $\lambda_0 + d\lambda = \lambda_1$ and $\lambda_1 - d\lambda = \lambda_0$ in order to use the Bennett Acceptance Ratio[S17] in estimating the free energy difference of a single $\lambda$ step, given by the following set of equations:

$$
\begin{aligned}
e^{\beta \Delta G_i} &= \frac{\langle f[\beta(U_0 - U_1 + c)]\rangle_1}{\langle f[\beta(U_1 - U_0 - c)]\rangle_0} e^{\beta c}, \\
c &= \Delta G_i + \frac{1}{\beta} \ln \frac{N_1}{N_0},
\end{aligned}
$$

where $f$ is the Fermi function $f(x) = 1/(1 + e^x)$, $\beta = 1/(k_B T)$, $U_1$ and $U_0$ are the energies for the forward and backward mutation, and $N_1$ and $N_0$ are the number of samples from each distribution of sampling in the forward and backward direction. In this case $N_0$ and $N_1$ are equal because the length of the forward and backward trajectories are the same, and conformations are sampled with identical rates. Summation across the entire lambda path gives the total free energy change, i.e. $\Delta G = \sum_i \Delta G_i$.

Each alchemical substitution is repeated three times starting from different initial structures chosen after equilibration. The values reported in the main text are the result of the average, and standard deviations were computed over these repetitions.

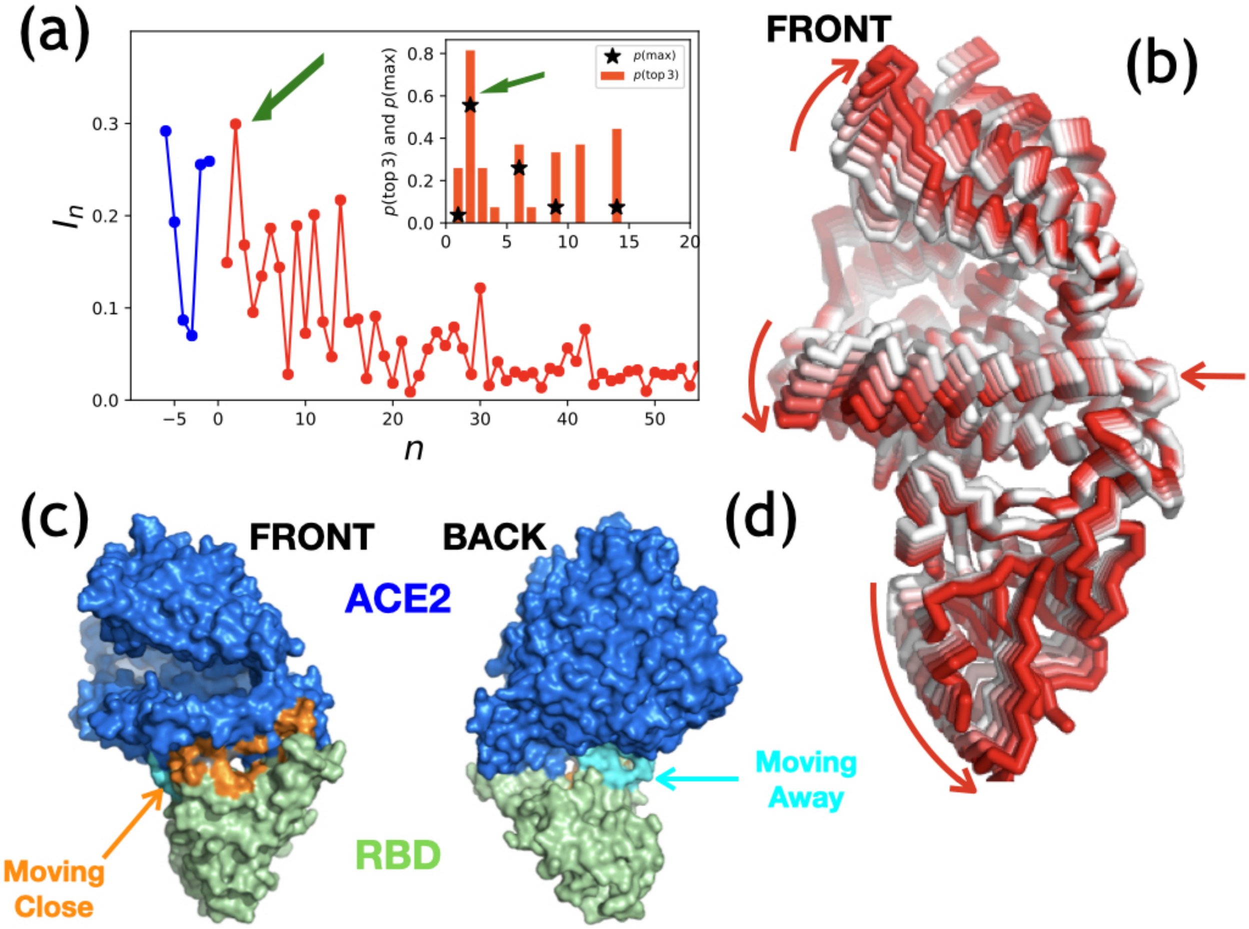


Figure S1: ENM analysis for SARS-CoV, PDBID: 2AJF.[S10] The figure is the same as Fig. 2 in the main text, but the analysis is performed on SARS-CoV. See Fig. 2 for details.

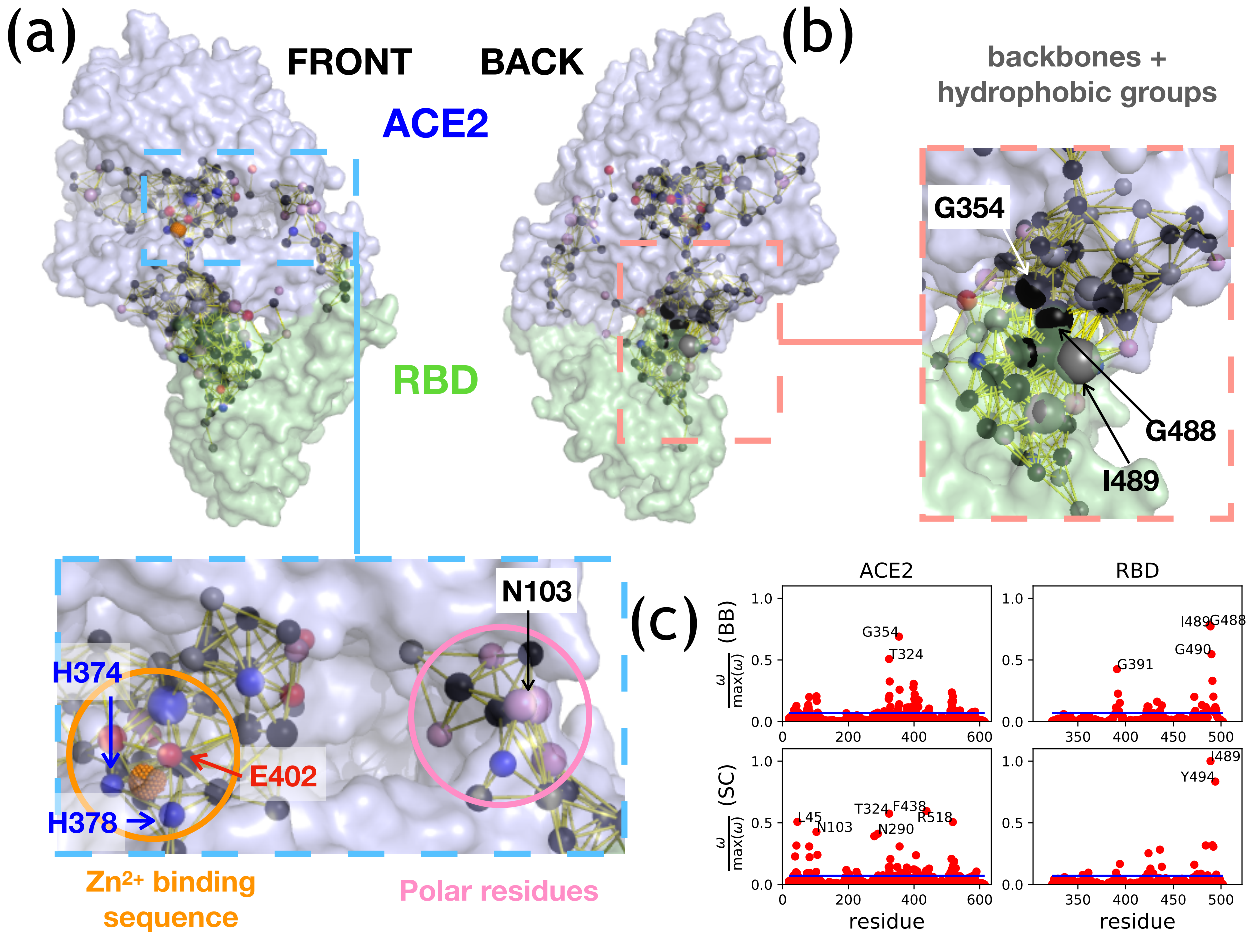


Figure S2: AWD for SARS-CoV. The figures are analogues of Fig. 3 in the main text except the results are for SARS-CoV. See Fig. 3 for details.

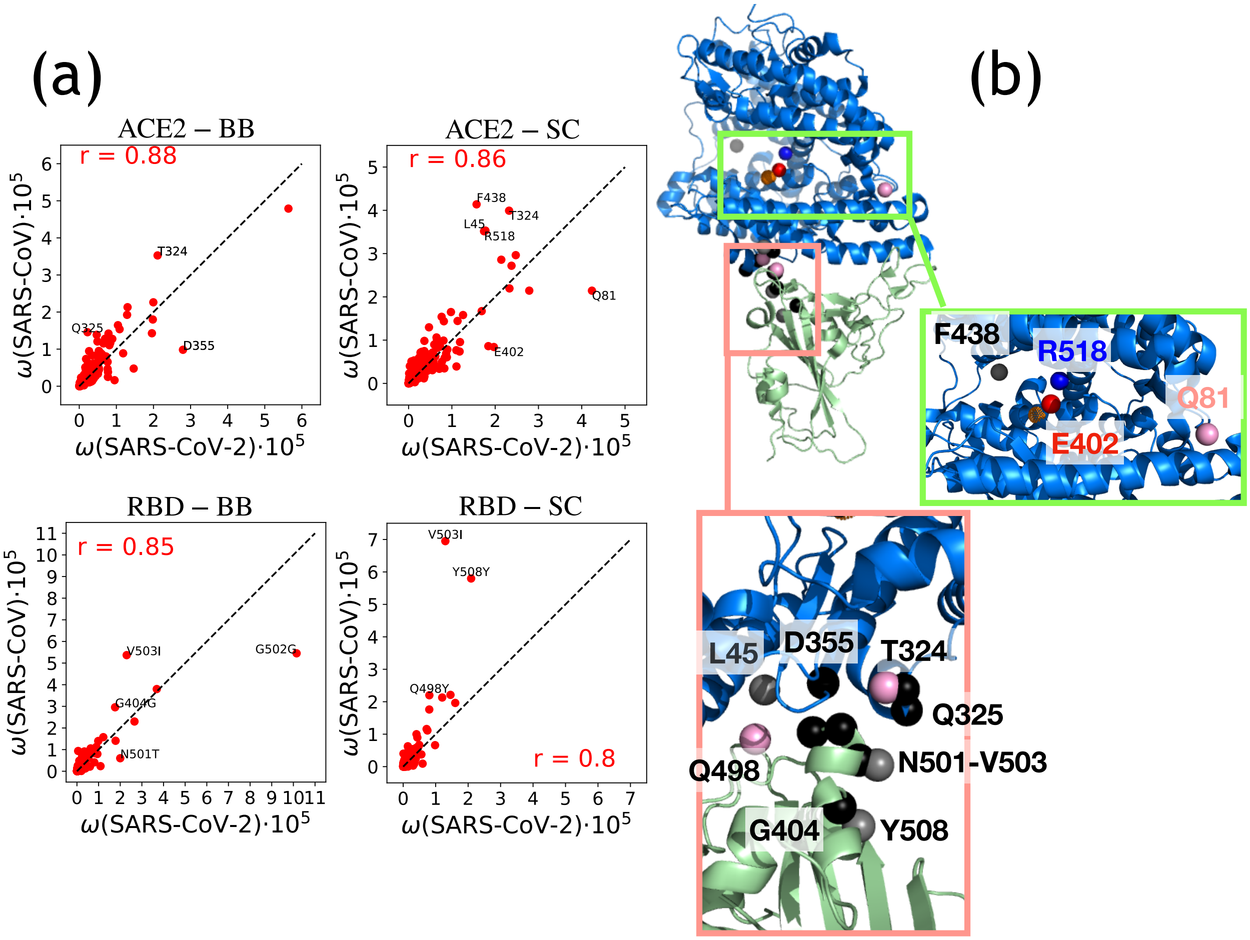


Figure S3: Comparison of SPM signal for SARS-CoV-2 and SARS-CoV. (a) Correlation of the SPM signal, with the values shown multiplied by $10^5$. The x-axis refers to SARS-CoV-2, the values for SARS-CoV are on the y-axis. The four panels show results for ACE2 backbone (top left) and side-chains (top right), and for backbone and side-chains of RBD (bottom left and right, respectively). For the RBD, each SARS-CoV-2 group is compared with the corresponding group of SARS-CoV after sequence alignment. The black line indicates the diagonal. The numbers in red are the values of the Pearson correlation coefficient. The groups for which the deviation between SARS-CoV-2 and SARS-CoV is larger than $10^5$ are labeled – the numbering refers to SARS-CoV-2. (b) Location of the groups (backbone or side chain) showing the largest deviation in the SPM signal between the two complexes. The green box highlights the section between subdomains I and II of the ACE2, including the active site, and the red box refers to the back of the complex, in the vicinity of G502. Residues are labeled according to the numbering in SARS-CoV-2.

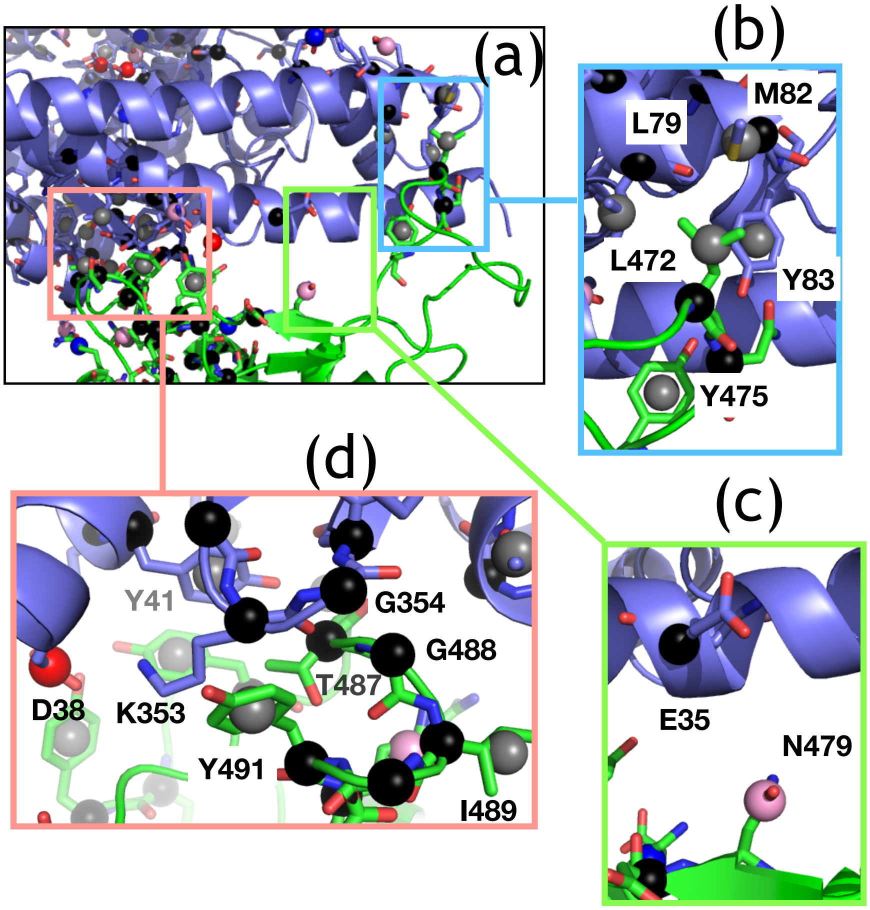


Figure S4: AWD at the interface of SARS-CoV. The analysis is the same as in Fig. 4. See the caption of Fig. 4 for details. The figures mirror those in Fig. 2 in the main text. See Fig. 2 for details.

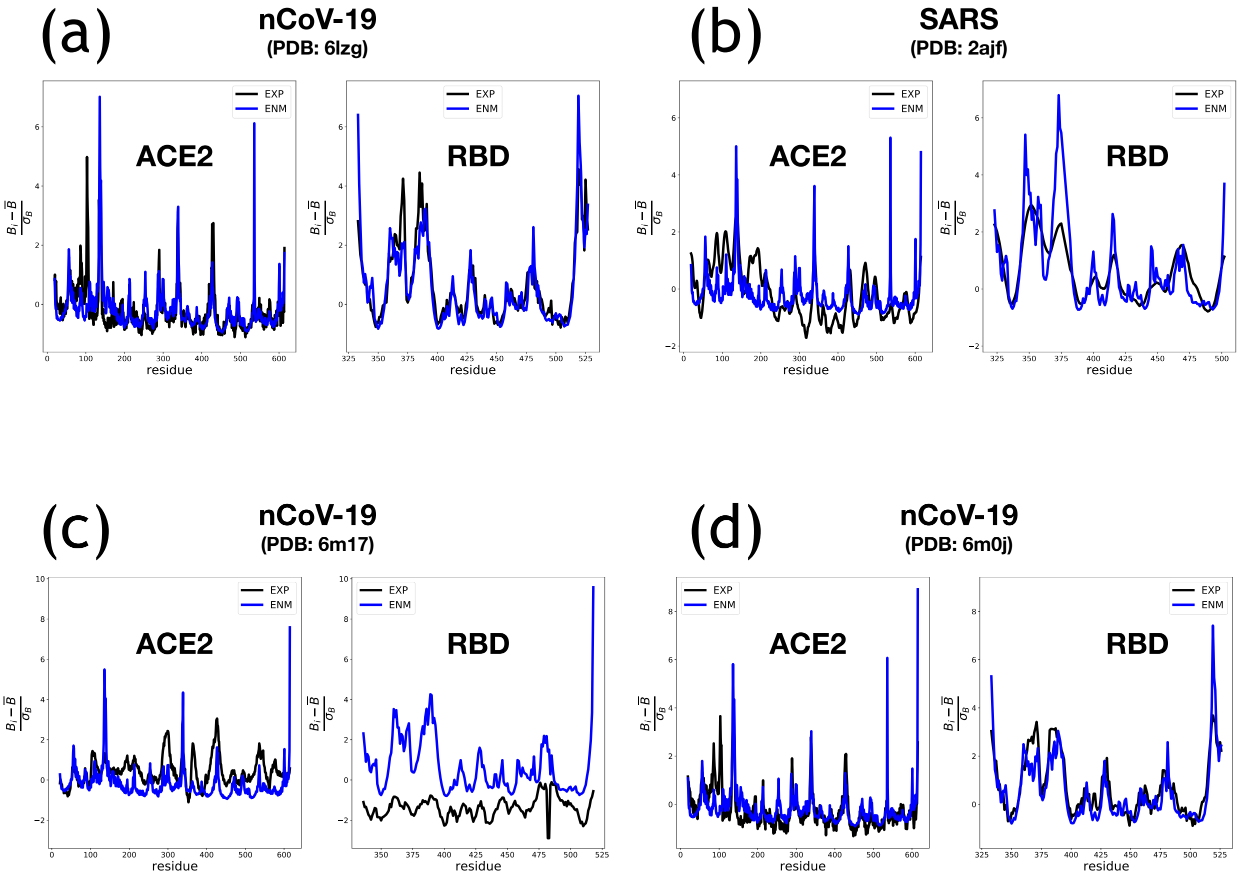


Figure S5: Comparison between experimental and calculated B-factors. Each panel is divided in 2, ACE2 and RBD. The plot shows the relative B-factors using Eq. S17. The black (blue) curve is from experiment (ENM). (a) SARS-Cov-2, PDBID: 6LZG,[S18] solved using X-ray diffraction at a resolution of 2.50 Å. (b) SARS-CoV, PDBID: 2AJF,[S10] solved via X-ray diffraction at a resolution of 2.90 Å. (c) SARS-Cov-2, PDBID: 6M17,[S12] considering only a subset of the ACE2 equivalent to the other structures. Solved using Cryo-EM at a resolution of 2.90 Å. (d) SARS-Cov-2, PDBID: 6M0J,[S19] solved via X-ray diffraction at a resolution of 2.45 Å.

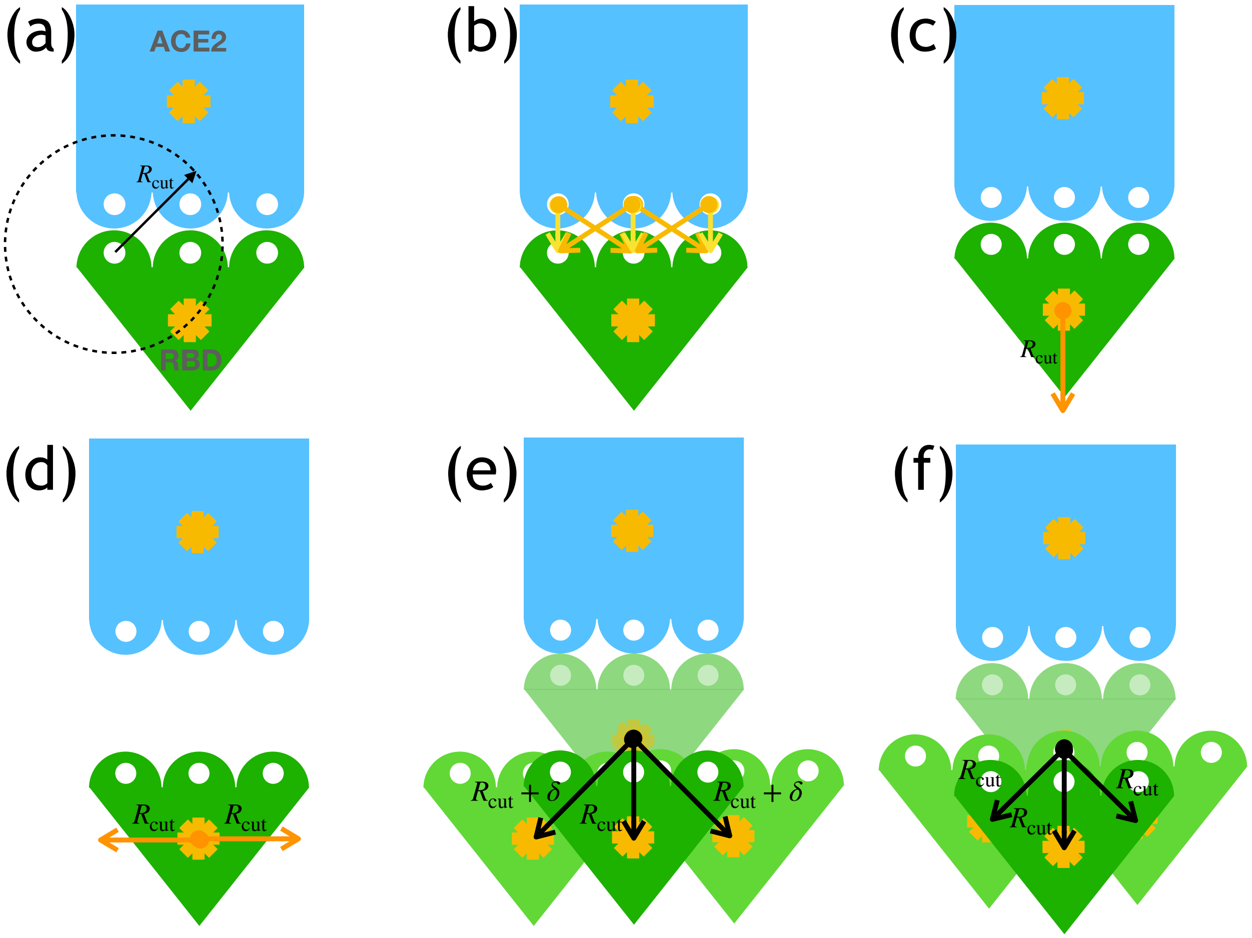


Figure S6: Construction of the ensemble of detached conformations. In all the panels, the blue and green pictures refer to the ACE2 and RBD, respectively. The white beads represent interaction sites at the interface, and the orange stars indicate the geometric center of the RBD and ACE2. (a) Bound conformation of the complex. The range of interaction (within $R_{\mathrm{cut}}$) for a selected site is represented with a black arrow and a black dashed line. Other sites follow the same rule. (b) Interaction pairs at the interface. All the interacting pairs at the interface are connected by yellow and orange arrows, which correspond to vectors connecting a bead from ACE2 and one from RBD within $R_{\mathrm{cut}}$ from each other. (c) The geometric center of the RBD is displaced by an amount $R_{\mathrm{cut}}$ away from ACE2, along a vector (orange in the picture) corresponding to the average of the vectors depicted in panel (b). (d) The "central" dissociated conformation is depicted. The "central" RBD is displaced by $R_{\mathrm{cut}}(i,j,k)$, where $(i,j,k)$ is a 3-dimensional vector of Cartesian components $i$, $j$, and $k$, with $i$, $j$, and $k \in \{-1,0,1\}$. Here only two arrows are shown in orange. (e) The displacement results in an ensemble of RBDs, here 3, which are displaced from the bound RBD (shown as a semi-transparent picture) by vectors of different directions and different length. (f) After the vectors are normalized to $R_{\mathrm{cut}}$, the ensemble of dissociated states is constructed, and so is the family of displacement vectors **D**-s, shown as black arrows.

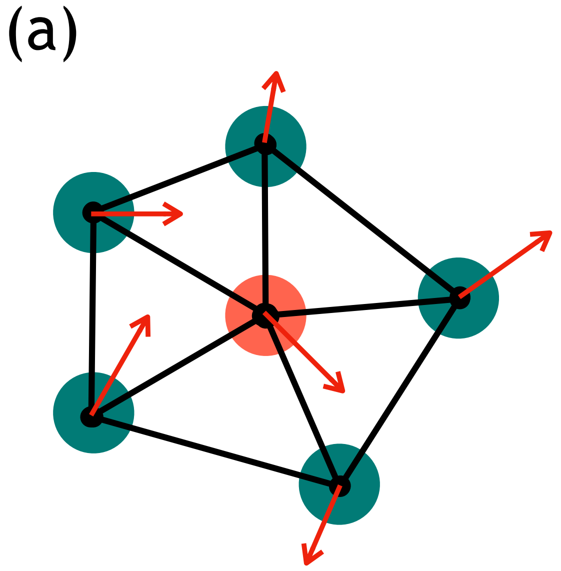


Figure S7: Structural Perturbation Method. (a) A network of interacting beads in blue and red; the interactions are shown as black lines. In the selected normal mode, each bead moves according to the depicted red arrows. (b) The network is perturbed by removing the interactions associated with the red bead.

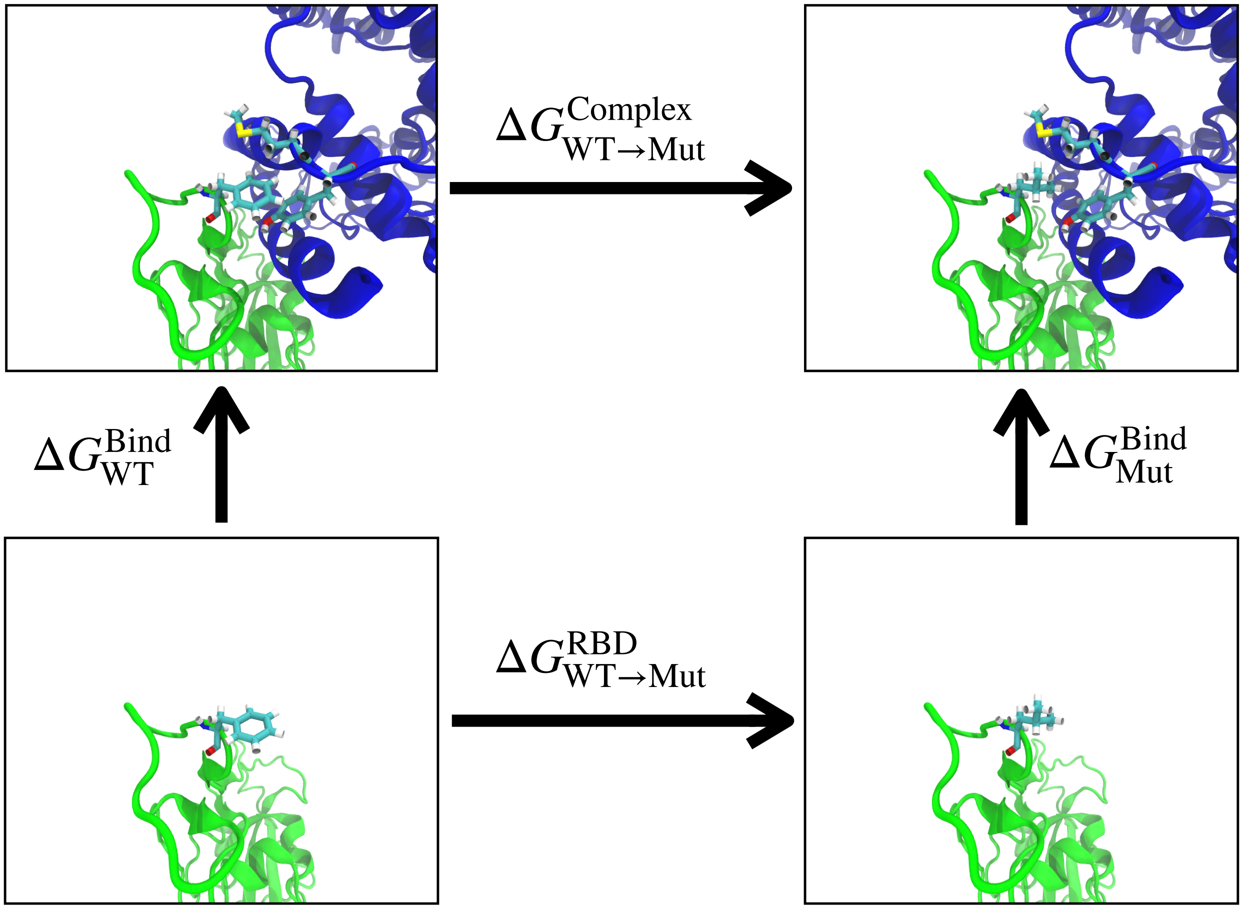


Figure S8: Scheme for calculation of relative binding free energies using alchemical substitutions. The four panels show the WT RBD bound to ACE2 (top left), the complex between the receptor and the mutant RBD (top right), the WT RBD in solution (bottom left), and the mutant RBD in the absence of ACE2 (bottom right). The sticks show F486 (WT) and L486 (mutant) on the RBD, together with M82 and Y83 on ACE2. Vertical arrows correspond to free energy differences associated with the formation of the complex, which can be measured experimentally. Horizontal arrows refer to free energy differences in the complex and with the RBD alone, which can be computed numerically via alchemical methods.

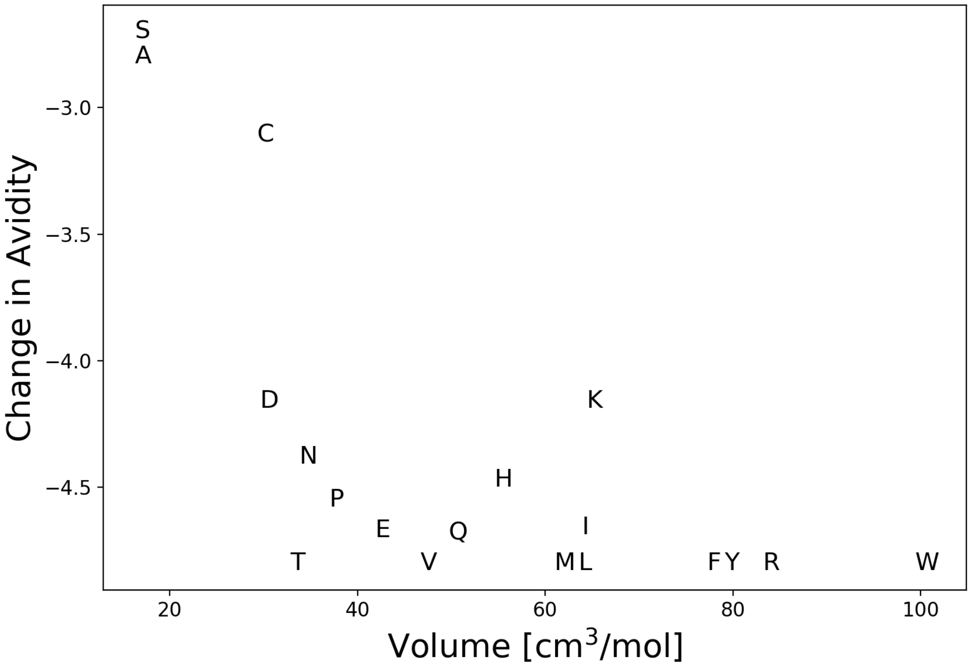


Figure S9: Correlation between size of the amino acid at location 502 in the RBD and the change in avidity for ACE2 of the G502X mutant compared to WT. The volumes on the x-axis are taken from Table 3, column 2 of Zamyatnin[S20] by subtracting the volume of glycine to the volume of each amino acid. The changes in avidity, taken from Starr *et al.*,[S21] report the difference of the base-10 logarithm of the apparent dissociation constant of the mutant (X, that is the residue indicate in the plot) and the wild type (glycine).

(S6) Oliphant, T. E. *A guide to NumPy*; Tregol Publishing, 2006.

(S7) van der Walt, S. J.; Colbert, S. C.; Gaël, V. The Numpy Array: A Structure for Efficient Numerical Computing. *Computing in Science & Engineering* **2011**, *13*, 22–30.

(S8) Virtanen, P. et al. SciPy 1.0: Fundamental Algorithms for Scientific Computing in Python. *Nature Methods* **2020**, *17*, 261–272.

(S9) Tehver, R.; Chen, J.; Thirumalai, D. Allostery Wiring Diagrams in the Transitions that Drive the GroEL Reaction Cycle. *Journal of Molecular Biology* **2009**, *387*, 390–406.

(S10) Li, F.; Li, W.; Farzan, M.; Harrison, S. C. Structure of SARS Coronavirus Spike Receptor-Binding Domain Complexed with Receptor. *Science* **2005**, *309*, 1864–1868.

(S11) Chipot, C.; Pohorille, A. *Free energy calculations*; Springer, 2007.

(S12) Yan, R.; Zhang, Y.; Li, Y.; Xia, L.; Guo, Y.; Zhou, Q. Structural basis for the recognition of SARS-CoV-2 by full-length human ACE2. *Science* **2020**, *367*, 1444–1448.

(S13) Vanommeslaeghe, K.; Hatcher, E.; Acharya, C.; Kundu, S.; Zhong, S.; Shim, J.; Darian, E.; Guvench, O.; Lopes, P.; Vorobyov, I.; Mackerell, A. D. CHARMM general force field: A force field for drug-like molecules compatible with the CHARMM all-atom additive biological force fields. *Journal of computational chemistry* **2009**, *31*, 671–NA.

(S14) Jorgensen, W. L.; Chandrasekhar, J.; Madura, J. D.; Impey, R. W.; Klein, M. L. Comparison of simple potential functions for simulating liquid water. *The Journal of chemical physics* **1983**, *79*, 926–935.

(S15) Phillips, J. C.; Braun, R.; Wang, W.; Gumbart, J.; Tajkhorshid, E.; Villa, E.; Chipot, C.; Skeel, R. D.; Kale, L.; Schulten, K. Scalable molecular dynamics with NAMD. *Journal of computational chemistry* **2005**, *26*, 1781–1802.

(S16) Humphrey, W.; Dalke, A.; Schulten, K. VMD: visual molecular dynamics. *Journal of molecular graphics* **1996**, *14*, 33–38.

(S17) Bennett, C. H. Efficient estimation of free energy differences from Monte Carlo data. *Journal of Computational Physics* **1976**, *22*, 245 – 268.

(S18) Wang, Q.; Zhang, Y.; Wu, L.; Niu, S.; Song, C.; Zhang, Z.; Lu, G.; Qiao, C.; Hu, Y.; Yuen, K.-Y.; Wang, Q.; Zhou, H.; Yan, J.; Qi, J. Structural and Functional Basis of SARS-CoV-2 Entry by Using Human ACE2. *Cell* **2020**, *181*, 894–904.e9.

(S19) Lan, J.; Ge, J.; Yu, J.; Shan, S.; Zhou, H.; Fan, S.; Zhang, Q.; Shi, X.; Wang, Q.; Zhang, L.; Wang, X. Structure of the SARS-CoV-2 spike receptor-binding domain bound to the ACE2 receptor. *Nature* **2020**, *581*, 215–220.

(S20) Zamyatnin, A. A. Amino Acid, Peptide, and Protein Volume in Solution. *Annual review of biophysics and bioengineering* **1984**, *13*, 145–165.

(S21) Starr, T. N.; Greaney, A. J.; Hilton, S. K.; Ellis, D.; Crawford, K. H.; Dingens, A. S.; Navarro, M. J.; Bowen, J. E.; Tortorici, M. A.; Walls, A. C.; King, N. P.; Veesler, D.; Bloom, J. D. Deep Mutational Scanning of SARS-CoV-2 Receptor Binding Domain Reveals Constraints on Folding and ACE2 Binding. *Cell* **2020**, *182*, 1295–1310.e20.
